# Evolution of a cytoplasmic determinant: evidence for the biochemical basis of functional evolution of a novel germ line regulator

**DOI:** 10.1101/2021.04.26.441385

**Authors:** Leo Blondel, Savandara Besse, Cassandra G. Extavour

**Affiliations:** Department of Molecular and Cellular Biology, Harvard University, Cambridge MA, USA; Department of Biochemistry and Molecular Medicine, Université de Montréal, Montréal Québec, Canada; Department of Organismic and Evolutionary Biology, Harvard University, Cambridge MA, USA

**Keywords:** *Oskar*, *vasa*. Drosophila, germ plasm, germ cell, LOTUS domain, RNA binding, Hidden Markov Models, Hymenoptera, Lepidoptera, Zygentoma

## Abstract

Germ line specification is essential in sexually reproducing organisms. Despite their critical role, the evolutionary history of the genes that specify animal germ cells is heterogeneous and dynamic. In many insects, the gene *oskar* is required for the specification of the germ line. However, the germ line role of *oskar* is thought to be a derived role resulting from co-option from an ancestral somatic role. To address how evolutionary changes in protein sequence could have led to changes in the function of Oskar protein that enabled it to regulate germ line specification, we searched for *oskar* orthologs in 1565 publicly available insect genomic and transcriptomic datasets. The earliest-diverging lineage in which we identified an *oskar* ortholog was the order Zygentoma (silverfish and firebrats), suggesting that *oskar* originated before the origin of winged insects. We noted some order-specific trends in *oskar* sequence evolution, including whole gene duplications, clade-specific losses, and rapid divergence. An alignment of all known 379 Oskar sequences revealed new highly conserved residues as candidates that promote dimerization of the LOTUS domain. Moreover, we identified regions of the OSK domain with conserved predicted RNA binding potential. Furthermore, we show that despite a low overall amino acid conservation, the LOTUS domain shows higher conservation of predicted secondary structure than the OSK domain. Finally, we suggest new key amino acids in the LOTUS domain that may be involved in the previously reported Oskar-Vasa physical interaction that is required for its germ line role.

## Introduction

With the evolution of obligate multicellularity, many organisms faced a challenge considered a major evolutionary transition: allocating only some cells (germ line) to pass on their genetic material to the next generation, relegating the remainder (soma) to death upon death of the organism (reviewed in (Kirk 2005)). This is soma-germ line differentiation, where only cells from the germ line will create the next generation (reviewed in (Kirk 2005)). While there are multiple mechanisms of germ cell specification, they can be grouped into two broad categories, induction or inheritance (reviewed in (Extavour and Akam 2003)). Under induction, cells respond to an external signal by adopting germ cell fate. Under the inheritance mechanism, maternally synthesized cytoplasmic molecules, collectively called germ plasm, are deposited in the oocyte and “inherited” by a subset of cells during early embryonic divisions. Cells inheriting these molecules commit to a germ line fate (reviewed in (Extavour and Akam 2003)).

The inheritance mechanism in insects that undergo metamorphosis (Holometabola) appears to have evolved by co-option of a key gene, *oskar*. *oskar* was first identified in forward genetic screens for axial patterning mutants in *Drosophila melanogaster* (Lehmann and Nüsslein-Volhard 1986). For the first 20 years following its discovery, *oskar* appeared to be restricted to Drosophilids (Clark, et al. 2007). Its later discovery in the mosquitoes *Aedes aegypti*, *Anopheles gambiae* and *Culex quinquefasciatus* (Juhn and James 2006; Juhn, et al. 2008) and the wasp *Nasonia vitripennis* (Lynch, et al. 2011) suggested the hypothesis that *oskar* emerged at the base of the Holometabola, and facilitated the evolution of germ plasm in these insects (Lynch, et al. 2011). However, our subsequent identification of *oskar* orthologs in the cricket *Gryllus bimaculatus* (Ewen-Campen, et al. 2012), and in many additional hemimetabolous insect species (Blondel, et al. 2020), demonstrated that *oskar* predates the Holometabola, and must be at least as old as the major radiation of insects (Misof, et al. 2014). Two secondary losses of *oskar* from insect genomes have also been reported, in the beetle *Tribolium castaneum* (Lynch, et al. 2011) and the honeybee *Apis mellifera* (Dearden, et al. 2006), and neither of these insects appear to use germ plasm to establish their germ lines (Nelson 1915; Nagy, et al. 1994; Dearden 2006; Schroder 2006). Whether *oskar* is ubiquitous across all insect orders, whether it is truly unique to insects, the evidence for or against potential losses or duplications of the *oskar* locus across insects, and the evolutionary dynamics of the locus, remain unknown.

*oskar* remains, to our knowledge, the only gene that has been experimentally demonstrated to be both necessary and sufficient to induce the formation of functional primordial germ cells (Kim-Ha, et al. 1991; Ephrussi and Lehmann 1992). Thus, in *D. melanogaster* (Lehmann and Nüsslein-Volhard 1986; Kim-Ha, et al. 1991; Ephrussi and Lehmann 1992) and potentially more broadly in holometabolous insects with germ plasm (Lynch, et al. 2011; Rafiqi, et al. 2020), *oskar* plays an essential germ line role. However, it is clear that *oskar*’s germ line function can evolve rapidly, as even within the genus *Drosophila*, *oskar* orthologs from different species cannot always substitute for each other (Webster, et al. 1994; Jones and Macdonald 2007). Moreover, the ancestral function of this gene may have been in the nervous system rather than the germ line (Ewen-Campen, et al. 2012). The current hypothesis is therefore that it was co-opted to play a key role in the acquisition of an inheritance-based germ line specification mechanism approximately 300 million years ago (Misof, et al. 2014), in the lineage leading to the Holometabola (Ewen-Campen, et al. 2012). Thus, the case of *oskar* offers an opportunity to study the evolution of protein function at multiple levels of biological organization, from the genesis of a novel protein, through to potential co-option events and the evolution of functional variation.

Neofunctionalization often correlates with a change in the fitness landscape of the protein sequence caused by novel biochemical constraints imposed by amino acid sequence changes (Sikosek, et al. 2012; Sikosek and Chan 2014). Such potential constraints may be revealed by analyzing the conservation of amino acids, their chemical properties, or structure at the secondary, tertiary or quaternary levels (Sikosek and Chan 2014). Oskar has two well-structured domains conserved across identified orthologs to date (Blondel, et al. 2020): an N-terminal Helix Turn Helix (HTH) domain termed LOTUS with potential RNA binding properties (Anantharaman, et al. 2010; Jeske, et al. 2015; Yang, et al. 2015; Jeske, et al. 2017), and a C-terminal GDSL-lipase-like domain called OSK (Jeske, et al. 2015; Yang, et al. 2015) (Figure 1). These two domains are linked by an unstructured highly variable interdomain sequence (Ahuja and Extavour 2014; Jeske, et al. 2015; Yang, et al. 2015). We previously showed that this domain structure is likely the result of a horizontal transfer event of a bacterial GDSL-lipase-like domain, followed by the fusion of this domain with a LOTUS domain in the host genome (Blondel, et al. 2020). Biochemical assays of the properties of the LOTUS and OSK domains provide some clues as to the molecular mechanisms that Oskar uses to assemble germ plasm in *D. melanogaster*. The LOTUS domain is capable of homodimerization (Jeske, et al. 2015; Jeske, et al. 2017), and directly binds and enhances the helicase activity of the ATP-dependent DEAD box helicase Vasa, a germ plasm component (Jeske, et al. 2017). The OSK domain resembles GDSL lipases in sequence (Jeske, et al. 2015; Yang, et al. 2015; Blondel, et al. 2020), but is predicted to lack enzymatic activity, as the conserved amino acid triad (S200 D202 H205) that defines the active site of these lipases is not conserved in OSK (Anantharaman, et al. 2010; Jeske, et al. 2015; Yang, et al. 2015). Instead, co-purification experiments suggest that OSK has RNA binding properties, consistent with its predicted basic surface residues (Jeske, et al. 2015; Yang, et al. 2015). Whether or how changes in the primary sequence of Oskar can explain the evolution of its molecular mechanism or tissue-specific function, remain unknown.

**Figure 1:**
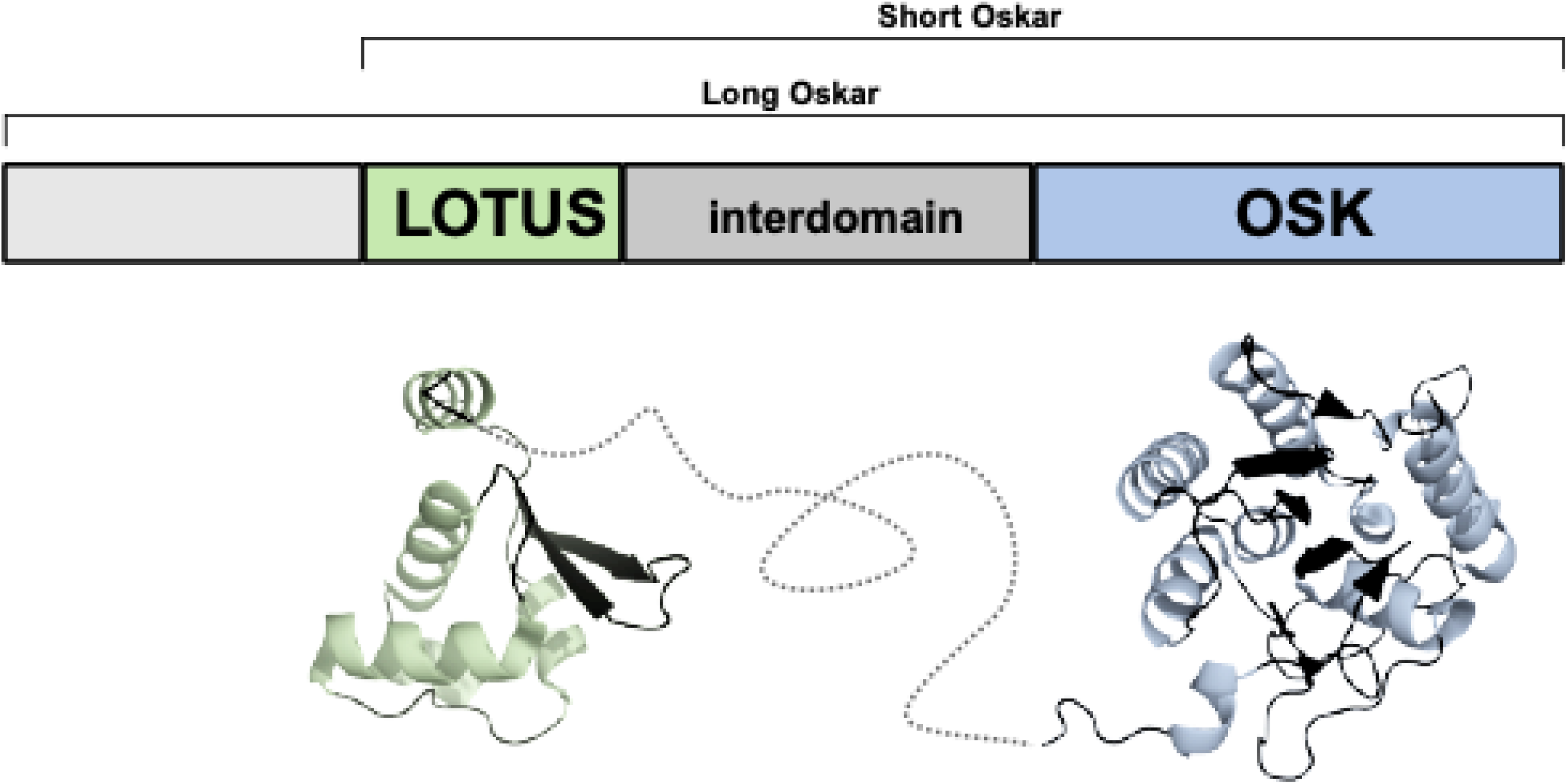
Overview of Oskar protein structure. The most common isoform of the Oskar protein, Short Oskar, is composed of two well-folded domains, LOTUS and OSK, separated by an interdomain sequence. A second isoform of the protein called Long Oskar is present in some Dipteran insects, which contains a 5’ domain as well as the three domains of Short Oskar. Below the schematic representation is a rendering of the previously reported solved structures for the LOTUS (PDBID: 5NT7) and OSK (PDBID: 5A4A) domains (Jeske, et al. 2015; Yang, et al. 2015) with a speculative rendering of the unfolded interdomain region shown with a dashed line

To date, sequences of approximately 100 *oskar* orthologs have been reported (Lynch, et al. 2011; Jeske, et al. 2015; Quan and Lynch 2016; Blondel, et al. 2020). However, the vast majority of these are from the Holometabola, and it is thus unclear whether analysis of these sequences alone would have sufficient power to allow extrapolation of conservation and divergence of putative biochemical properties across insects broadly speaking. Multiple hypotheses as to the molecular mechanistic function of particular amino acids in the LOTUS and OSK domains in *D. melanogaster* have been proposed (Jeske, et al. 2015; Yang, et al. 2015; Jeske, et al. 2017), but without sufficient taxon sampling, the potential relevance of these mechanisms to *oskar*’s evolution and function in other insects is unclear.

Here we address these outstanding questions by applying a rigorous bioinformatic pipeline to generate the most complete collection of *oskar* sequences to date. By analyzing 1862 Pancrustacean genomes and transcriptomes, we show that *oskar* likely first arose at least 400 million years ago, before the advent of winged insects (Pterygota). We find that the *oskar* locus has been lost independently in some insect orders, including near-total absence from the order Hemiptera, and clarify that the absence of *oskar* from the *Bombyx mori* and *Tribolium castaneum* genomes (discussed in Quan and Lynch 2016) does not reflect a general absence of *oskar* from Lepidoptera or Coleoptera. By comparing Oskar sequences in a phylogenetic context, we reveal that distinct biophysical properties of Oskar are associated with Hemimetabola and Holometabola. We use these observations to propose testable hypotheses regarding the putative biochemical basis of evolutionary change in Oskar function across insects.

## Results

### HMM-based discovery pipeline yields hundreds of novel oskar orthologs

We wished to study the evolution of the *oskar* gene sequence as comprehensively as possible across all insects. To expand our previous collection of nearly 100 orthologous sequences (Blondel, et al. 2020), we designed a new bioinformatics pipeline to scan and search for *oskar* orthologs across all 1565 NCBI insect transcriptomes and genomes that were publicly available at the time of analysis (Supplementary Table S1; Figure 2; see Methods: *Genome and transcriptome pre-processing* for NCBI accession numbers and additional information). First, we used the HMMER tool suite to build HMM models for each of the LOTUS and OSK domains, using our previously generated multiple sequence alignments (MSA) (Blondel, et al. 2020). We subjected genomes to *in silico* gene model inference using Augustus (Stanke, et al. 2006). We translated the resulting predicted transcripts, as well as the predicted transcripts from RNA-seq datasets, in all six frames. We then scanned the resulting protein sequences for the presence of LOTUS and OSK domains using the aforementioned HMM models. Sequences were designated as *oskar* orthologs based on the same criteria as in our previous study (Blondel, et al. 2020), namely, sequences containing both a LOTUS and an OSK domain (Jeske, et al. 2015), separated by a variable interdomain region. We then aligned all sequences using *hmmalign* and the HMM derived from our previously published full length Oskar alignment (Blondel, et al. 2020), and manually curated sequence duplicates and sequences that did not align correctly.

**Figure 2:**
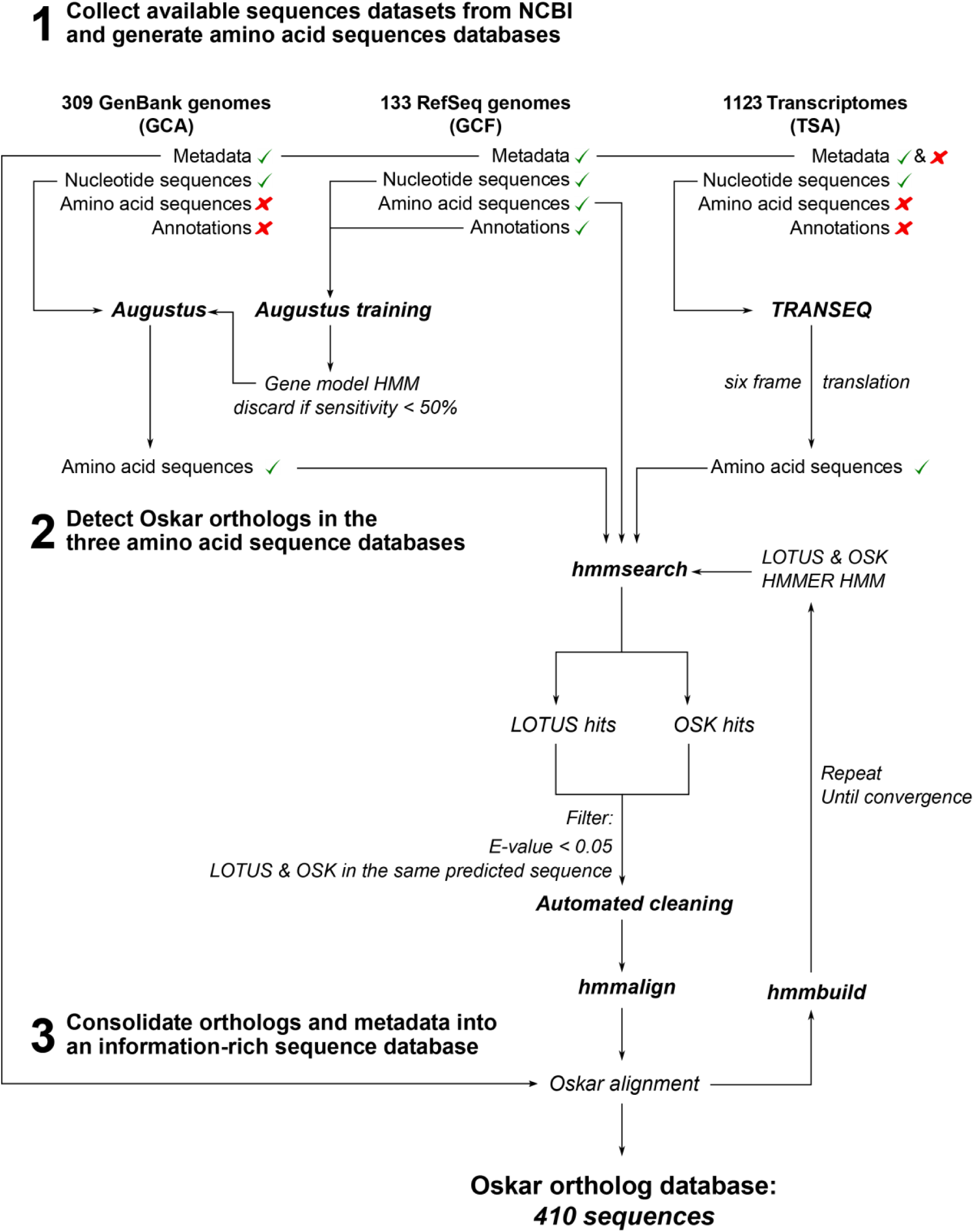
Schematic presentation of the *oskar* ortholog detection pipeline. Sequences were collected automatically from the three NCBI databases, GenBank (GCA), RefSeq (GCF) and Transcriptome Shotgun Assembly Database (TSA). RefSeq genomes were used to generate Augustus gene model HMMs, which were used to annotate and predict proteins in the non-annotated genomes obtained from GenBank. Transcripts from the TSA database were 6-frame translated using TRANSEQ. Amino acid sequences were consolidated into three protein databases. *hmmsearch* from the HMMER tool suite was used to search for LOTUS and OSK hits in those sequences. Sequences with hits for both the LOTUS and OSK domains with an E-value <0.05 were annotated as *oskar* sequences. Sequences were then cleaned to remove duplicates (sequences with <80% sequence similarity coming from the same organism). The resulting sequences were aligned using *hmmalign*, and the process was repeated until no new sequences were identified. Finally, the sequences were consolidated with the dataset metadata into the *oskar* ortholog database that was used for all subsequent analyses.

With these methods, we recovered a total of 379 unique *oskar* sequences from 350 unique species. To our knowledge, this comprises the largest collection of *oskar* orthologs described to date. To determine if *oskar* orthologs might predate Insecta, we applied the discovery pipeline to all 31 genomes and 266 transcriptomes of non-insect pancrustaceans available at the time of analysis (see Methods: *Genomes and transcriptomes preprocessing* for complete list). However, we did not recover any non-insect sequences meeting our criteria for *oskar* orthologs (Figure 3), strongly suggesting that *oskar* is restricted to the insect lineage (Lynch, et al. 2011; Ahuja and Extavour 2014).

**Figure 3:**
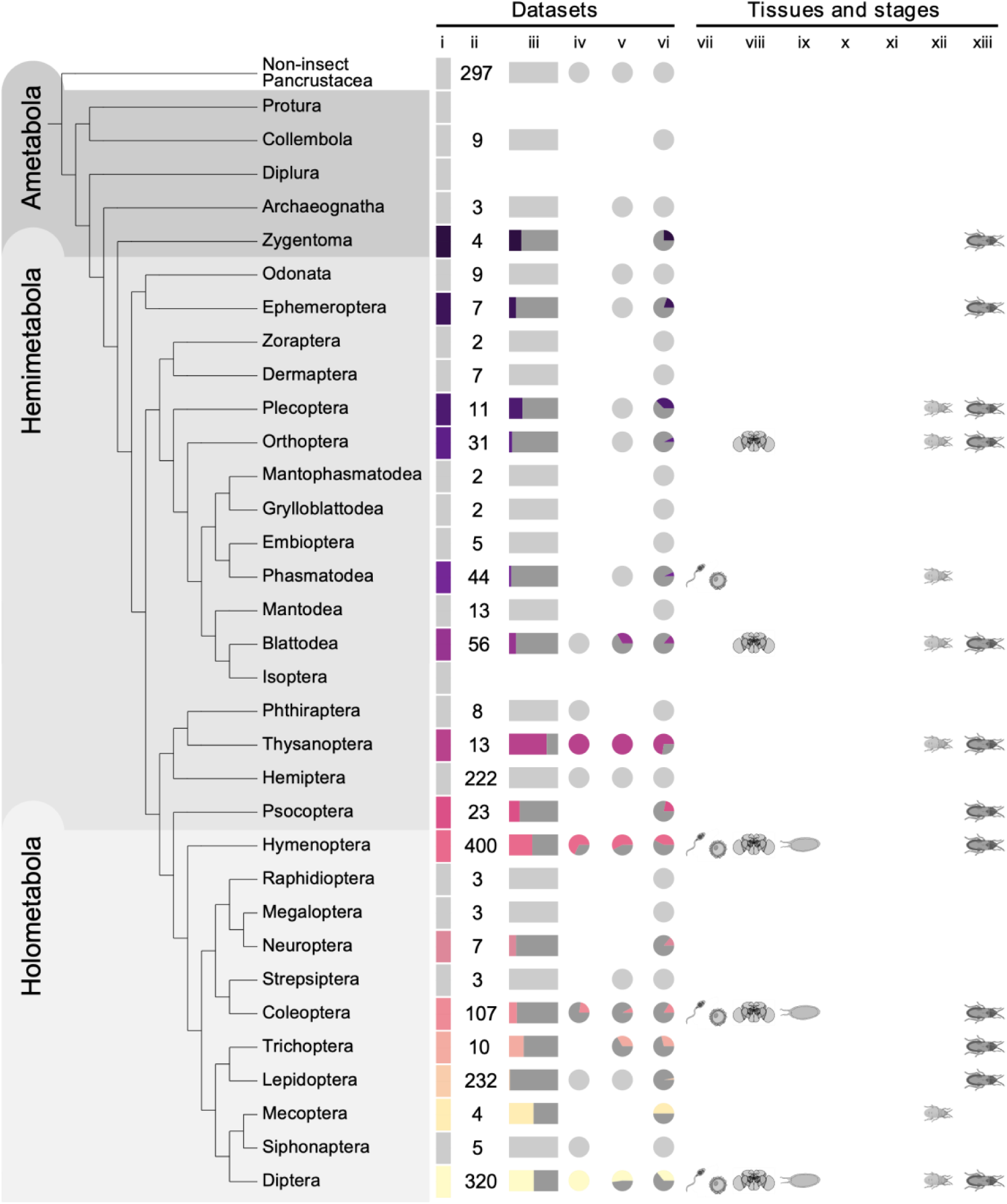
Summary of *oskar* distribution and expression in insects. Phylogeny from (Misof, et al. 2014). Symbols in order from left to right: (i) vertical rectangles: grey: no *oskar* ortholog was identified in this order. Color (unique for each order): at least one *oskar* ortholog was identified in this order. (ii) number of datasets searched. (iii) horizontal rectangles: proportion of searched datasets in which an *oskar* ortholog was identified. (iv) pie chart: proportion of *oskar* sequences identified in RefSeq (GCF) datasets. (v) pie chart: proportion of *oskar* sequences identified in GenBank (GCA) datasets. (vi) pie chart: proportion of *oskar* sequences identified in Transcriptome Shotgun Assembly Database (TSA) datasets. (vii) *oskar* sequences identified in tissue related to germ line (transcriptomes derived from reproductive organs, eggs or embryos). (viii) *oskar* sequences identified in tissue related to the brain (transcriptomes derived from brain or head). (ix) *oskar* sequences identified in an egg stage transcriptome. (x) *oskar* sequences identified in a larval stage transcriptome. (xi) *oskar* sequences identified in a pupal stage transcriptome. (xii) *oskar* sequences identified in a nymphal or juvenile stage transcriptome. (xiii) *oskar* sequences identified in an adult transcriptome. All numbers represented graphically here are in Supplementary Table 1. No datasets were available for Protura, Diplura or Isoptera at the time of analysis.

We found that 58.65% of RefSeq genomes (78/133), 30.42% of GenBank genomes (94/309), and 21.19% of transcriptomes (238/1123) analyzed contained predicted *oskar* orthologs (Supplementary Table S1 and Supplementary Figure S1a). Given that detection of putative orthologs is highly dependent on the quality of the genome assembly and annotation, we asked whether there were differences in the assembly statistics of genomes with and without predicted *oskar* orthologs. We observed a significant difference in N50, L50, number of contigs and number of scaffolds between genomes lacking *oskar* hits and those where *oskar* was identified (Mann-Whitney U test p-value < 0.05). Genomes where we did not find *oskar* showed a significantly higher mean/median contig and scaffold count, smaller contig and scaffold N50 length, larger contig and scaffold L50, and more contigs or scaffolds per genome length, than genomes where we detected an *oskar* ortholog (Mann Whitney U test p<0.05; Supplementary Figure S2; Supplementary Table S2).

### oskar predates the divergence of Ametabola and other insects

We identified *oskar* orthologs in 15 of the 29 generally recognized (Misof, et al. 2014) insect orders, including eight holometabolous orders, six hemimetabolous orders, and one ametabolous order (Figure 3). This result is consistent with our previous proposals that *oskar* predates the origins of the Holometabola (Ewen-Campen, et al. 2012; Blondel, et al. 2020). The novel finding of an *oskar* ortholog from the silverfish *Atelura formicaria* (Zygentoma) allows us to date back the origin of *oskar* further than previous analyses, to at least 420 million years ago (Misof, et al. 2014), before the divergence of Ametabola from the remaining insect lineages.

We then explored the distribution of *oskar* sequences across insect phylogeny. Interestingly, we identified multiple lineages where *oskar* appeared to have been lost independently, including confirming the previously reported (Lynch, et al. 2011) losses from the genomes of the red flour beetle *Tribolium castaneum*, the honeybee *Apis mellifera*, and the silk moth *Bombyx mori* (Figure 3). Notably, within Lepidoptera we identified *oskar* orthologs in only four species, despite the fact that we searched 232 available lepidopteran sequence datasets, including 17 well-annotated RefSeq genomes and 135 transcriptomes (Figure 3 and Supplementary Figure S3). In principle, this apparent widespread absence of *oskar* in Lepidoptera could be due to unusually rapid evolution of the *oskar* sequence in this lineage, which might render lepidopteran *oskar* orthologs undetectable by our methods. However, we note that the only four lepidopteran orthologs we detected all belonged to species of the basally branching *Adelidae* and *Palaephatidae* families. We therefore favor the interpretation that *oskar* was lost from a last common ancestor of *Meessiidae* and *Palaphaetidae*, approximately 180 million years ago, with the consequence that the majority of extant lepidopteran lineages lack an *oskar* ortholog (Supplementary Figure S3) (Mitter, et al. 2017; Kawahara, et al. 2019).

The Hemiptera also appear to have lost *oskar*, based on our analysis of the 222 datasets available for this clade, including 12 RefSeq genomes and 192 transcriptomes. However, we did identify an *oskar* ortholog in the Thysanoptera, which is a hemipteran sister group (Misof, et al. 2014). Finally, we identified *oskar* orthologs in only four of the 11 orders of the Polyneoptera for which data were available. With the exception of Mantodea (13 transcriptomes), the four orders with detectable *oskar* sequences all had more than ten available sequence datasets (Plecoptera: three genomes and eight transcriptomes; Orthoptera: three genomes and 28 transcriptomes; Phasmatodea: 13 genomes and 31 transcriptomes; Blattodea: five genomes and 51 transcriptomes). The remaining orders had fewer than eight datasets each available for analysis (Figure 3; Supplementary Table S1), which could account for the apparent paucity of *oskar* genes in this group. However, we cannot rule out the possibility that *oskar* in the Polyneoptera may have diverged beyond our ability to detect it, or that it may have been lost multiple times, as observed for multiple holometabolous orders.

As well as multiple convergent losses of *oskar*, we also uncovered evidence for independent instances of duplication of the *oskar* locus. We defined a putative duplication instance as two or more *oskar* sequences (possessing both a LOTUS and OSK domain as per our definition) in the same species that shared less than 80% sequence similarity. All of these events were detected within the Hymenoptera. We therefore performed a phylogenetic analysis of the hymenopteran sequences to test the hypothesis that these were the result of duplication events (Figure 4; Supplementary Figure S4). Our analysis of hymenopteran *oskar* sequences recovered previously published hymenopteran phylogenetic relationships (Peters, et al. 2017). We found that *oskar* was duplicated in the four Figitidae species studied, a family of parasitoid wasps. Moreover, one out of ten examined Cynipidae species, as well as the only Ceraphronidae species examined, also harbored a duplicated *oskar* sequence. Multiple *oskar* duplications were also identified in the Chalcidoid wasps, notably in the Mymaridae (all three species studied), the Eupelmidae (two out of three species), the Aphelinidae (both species) and the Pteromalidae (one out of 17 species). Finally, we identified two additional apparently independent duplication events in the Aculeata, one in the wasp *Polistes fuscatus* (of 29 Vespidae, including three additional *Polistes* species, two with RefSeq genomes (*P. canadensis* and *P. dominula*) in which *oskar* was identified in single copy), and one in the red imported fire ant *Solenopsis invicta* (of 41 Formicidae species, including the congeneric *S. fugax*, with a GenBank genome in which *oskar* was identified in single copy).

**Figure 4:**
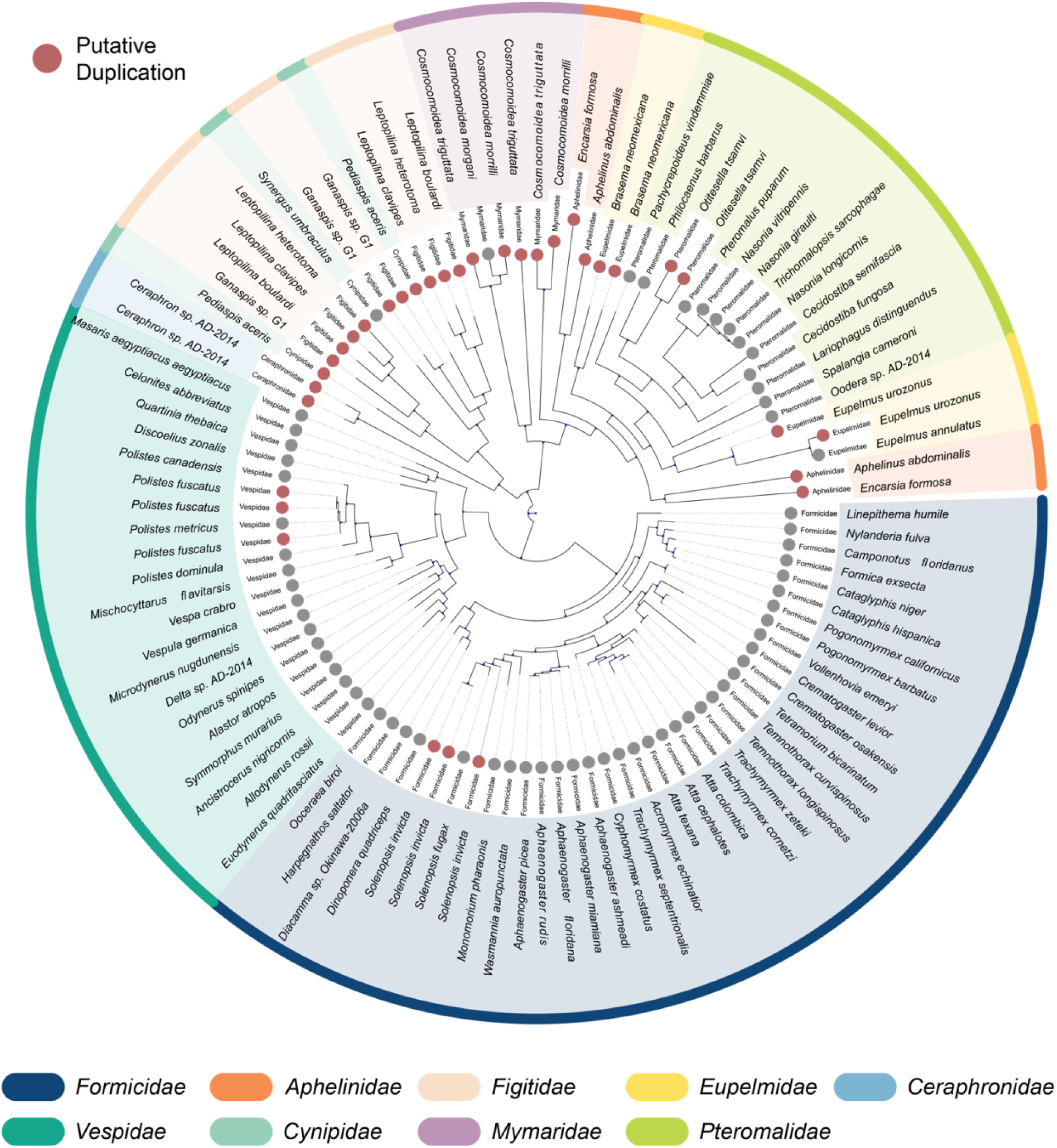
Phylogenetic reconstruction of hymenopteran Oskar sequences. Phylogenetic tree inferred using RaxML with 100 bootstraps. Each leaf represents an Oskar ortholog. Gray circles: only one Oskar sequence was identified. Red circles: putatively duplicated Oskar sequences identified (sequence similarity <80%). Only families which contained a putative duplication are shown here; see Supplementary Figure S4 for the results of our *oskar* search in the context of a more complete hymenopteran phylogeny.

### Evidence for oskar expression in multiple somatic tissues

In studied insects to date, *oskar* is expressed and required in one or both of the germ line (Juhn and James 2006; Juhn, et al. 2008; Lynch, et al. 2011; Lehmann 2016) or the nervous system (Ewen-Campen, et al. 2012; Xu, et al. 2013). We asked whether these expression patterns could be detected in the insects studied here. To this end, we downloaded all available metadata for the transcriptomes analyzed here, to obtain information on the source tissues and developmental stages. We obtained these data for 371 out of the 1123 transcriptomes in our analysis, including both holometabolous and hemimetabolous orders (see Methods: *TSA metadata parsing and curation*). To first explore the distribution of *oskar* expression in the brain and the germ line, we binned the different tissues reported in the metadata into two categories, brain or germ line. This was done independently of the developmental stage (if that information was included in the metadata) by creating a mapping table and checking the extracted tissues against this table (Supplementary Table S3 at GitHub repository ***TableS3_germline_brain_table.csv***). We then cross referenced our orthology detection with these metadata. We found evidence for *oskar* expression in the germ line of four orders (Phasmatodea, Hymenoptera, Coleoptera and Diptera), and in the brain of five orders (Orthoptera, Blattodea, Hymenoptera, Coleoptera, Diptera) (see Methods: *TSA metadata parsing and curation* for details on keyword extractions). In addition, we found evidence of *oskar* expression in several somatic tissues not previously implicated in studies of *oskar* expression and function. These tissues included the midgut (*Polistes fuscatus, Sitophilus oryzae*), fat body (*Polistes fuscatus, Arachnocampa luminosa*), salivary gland (*Culex tarsalis, Anopheles aquasalis, Leptinotarsa decemlineata*), venom gland (*Culicoides sonorensis, Fopius arisanus*), and silk gland (*Bactrocera cucurbitae*) (Supplementary Figure S5). In terms of developmental stage, only holometabolous insects appeared to express *oskar* during embryonic, larval or nymphal stages; for all other insects, *oskar* was detected in transcriptomes derived from adults (Figure 3). However, it is important to note that for most species, transcriptomes were available only from adult tissues, rather than from a full range of developmental stages (Supplementary Figure S5). We therefore cannot rule out the possibility that *oskar* expression at pre-adult stages is also a feature of multiple Hemimetabola. Indeed, we previously reported that *oskar* is expressed and required in the embryonic nervous system of a cricket, a hemimetabolous insect (Ewen-Campen, et al. 2012).

### The Long Oskar domain is an evolutionary novelty specific to a subset of Diptera

*D. melanogaster* has two isoforms of Oskar (Markussen, et al. 1995): Short Oskar, containing the LOTUS, OSK and interdomain regions, and Long Oskar, containing all three domains of Short Oskar as well as an additional 5’ domain (Supplementary Figure S7). It was previously reported that Long Oskar was absent from *N. vitripennis*, *C. pipiens* and *G. bimaculatus* (Lynch, et al. 2011; Ewen-Campen, et al. 2012), and within our alignment of Oskar sequences we could only detect the Long Oskar isoform within Diptera. Therefore, using our dataset, we asked when these two isoforms had evolved. We selected the dipteran sequences from our Oskar alignment and then grouped the sequences by family. We plotted the amino acid occupancy at each alignment position (Supplementary Figure S7), and found that Long Oskar predates the Drosophilids, being identified as early as the *Pinpunculidae* (Supplementary Figure S7). Moreover, following the evolution of the Long Oskar isoform, the Long Oskar domain was retained in all families except for the *Glossinidae* and *Scathophagidae*. However, given that we identified only eight and two Oskar sequences for these families respectively, we cannot eliminate the possibility that apparent absence of the Long Oskar domain in these groups reflects our small sample size, rather than true evolutionary loss.

### The LOTUS and OSK domains evolved differently between hemimetabolous and holometabolous insects

The fact that an *oskar-*dependent germ plasm mode of germ line specification mechanism has been identified only in holometabolous insects suggests that *oskar* may have been co-opted in this clade for this function (Ewen-Campen, et al. 2012). Under this hypothesis, evolution of the *oskar* sequence in the lineage leading to the Holometabola may have changed the physico-chemical properties of Oskar protein, such that it acquired germ plasm nucleation abilities in these insects. To test this hypothesis, we asked whether there were particular sequence features associated with Oskar proteins from holometabolous insects, in which Oskar can assemble germ plasm, and hemimetabolous insects, which lack germ plasm. In particular, we assessed the differential conservation of amino acids at particular positions across Oskar and asked if these might be predicted to change the physico-chemical properties of Oskar in specific ways that could potentially be relevant to germ plasm nucleation. We used the Valdar score (Valdar 2002) as the main conservation indicator for this study (see GitHub file ***scores.csv***), as this metric accounts not only for transition probabilities, stereochemical properties and amino acid frequency gaps, but also for the availability of sequence diversity in the dataset. It computes a weighted score, where sequences from less well-represented clades contribute proportionally more to the score than sequences from overrepresented clades. Due to the highly unbalanced availability of genomic and transcriptomic data between hemimetabolous and holometabolous sequences (Supplementary Table S1; Figure 3) the choice of a weighted score was necessary to avoid biasing the results towards insect orders such as Diptera or Hymenoptera. To study the difference between hemimetabolous and holometabolous sequences, we did not use the Valdar score directly, but instead computed the conservation ratio between both groups for each position, which we call the Conservation bias (See Methods: Computation of the Conservation Bias). We plotted the conservation bias on the solved three-dimensional crystal structure of the *D. melanogaster* LOTUS and OSK domains (Jeske, et al. 2015; Yang, et al. 2015) to ask whether specific functionally relevant structures showed phylogenetic or other patterns of residue conservation (Figure 5).

**Figure 5:**
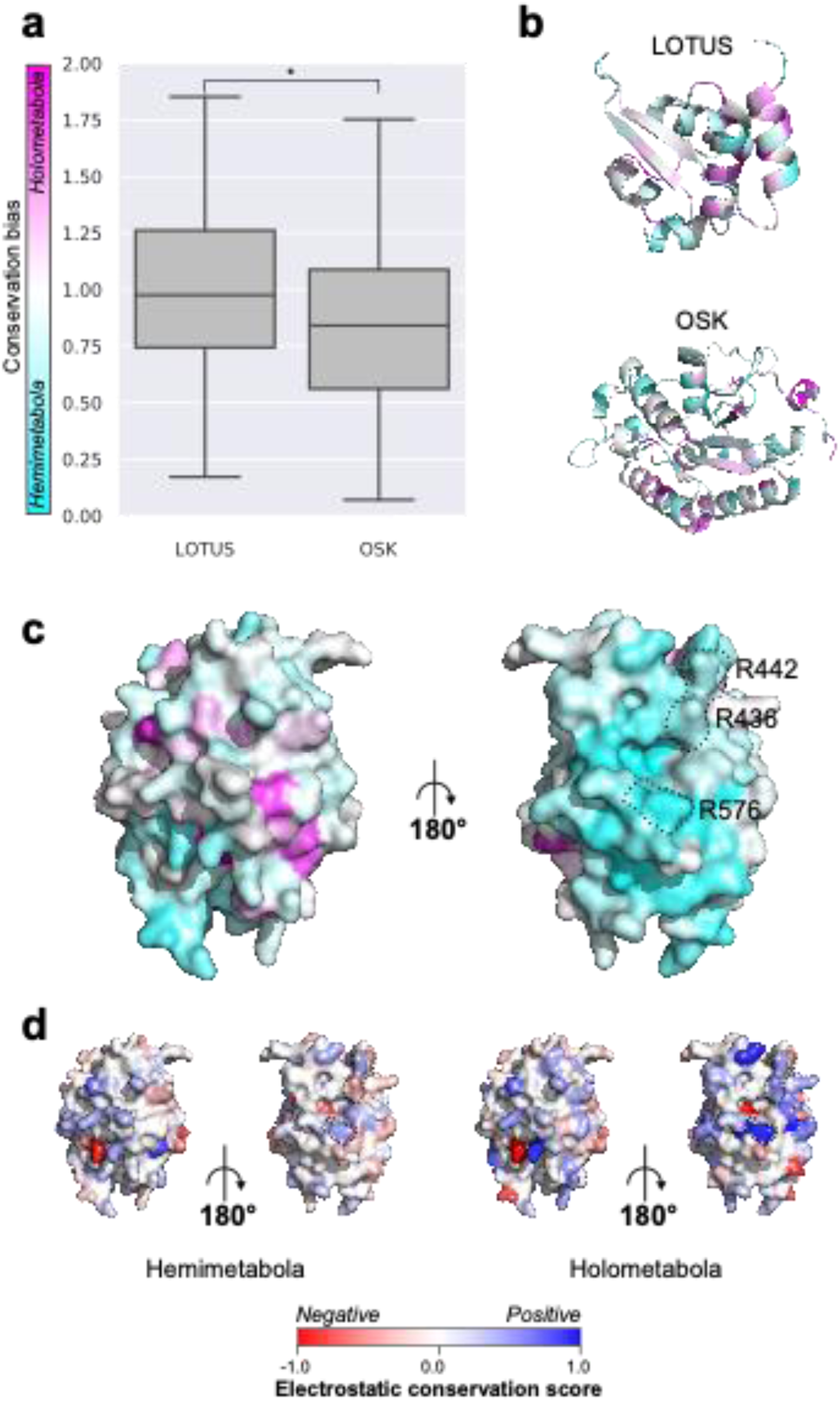
Differential conservation of amino acids between hemimetabolous and holometabolous Oskar sequences. **(a)** Box plot showing the conservation bias for each of the LOTUS and OSK domains between hemimetabolous and holometabolous Oskar sequences. Statistical difference was tested using a Mann Whitney U test (p <0.05). **(b)** Ribbon diagram of LOTUS (PDBID: 5NT7) and OSK (PDBID: 5A4A) domain structures, where each amino acid is colored by conservation bias on the color scale shown in **(a)**. **(c, d)** Protein surface representation of the OSK domain (PDBID: 5A4A) from two different angles. Black dashed lines indicate the three amino acids reported previously to be necessary for OSK binding to RNA in *D. melanogaster* (Jeske, et al. 2015; Yang, et al. 2015). **(c)** Amino acids colored by conservation bias on the color scale shown in **(a)**. Cyan: amino acids more highly conserved in hemimetabolous sequences; magenta: amino acids more highly conserved in holometabolous sequences. **(d)** Amino acids colored by electrostatic conservation score. Left: hemimetabolous sequences; right: holometabolous sequences.

First, we asked if the overall conservation score of the domains was different between holometabolous and hemimetabolous sequences. We observed that the conservation bias for the LOTUS domain was centered around a mean of 1.00, indicating that both Holometabola and Hemimetabola displayed a similar conservation of the LOTUS domain (Figure 5a). For the OSK domain however, the conservation bias was centered around 0.84, indicating that the hemimetabolous sequences displayed a higher level of conservation compared to holometabolous sequences (Figure 5a). We then looked at the conservation bias scores *in-situ* on the LOTUS domain structure. We asked if the amino acids of the ꞵ sheets of the LOTUS domain thought to be involved in dimerization of the protein (Jeske, et al. 2015; Yang, et al. 2015) displayed conservation bias. Both ꞵ sheets had an overall even bias (mean: 1.03 and 1.05 for ꞵ1 and ꞵ2 respectively) between both groups (Figure 5b). Second, as we had observed that hemimetabolous OSK was more conserved overall than holometabolous OSK, we asked if there were any clear patterns of conservation bias in specific regions of the structure (Figure 5a and b). We found that some of the secondary structures within OSK showed a differential conservation (α2: 0.54, α6: 0.42, ꞵ2: 0.52), whereas other structures were within less than 0.1 of the median value for OSK. Moreover, we observed a large pocket of amino acids showing a conservation bias towards hemimetabolous sequences located on the surface of OSK (Figure 5c). This particular area contains the previously reported important amino acids for the RNA binding function of OSK (Jeske, et al. 2015; Yang, et al. 2015) namely, R442, R436 and R576. The electrostatic properties at those positions were conserved in the holometabolous sequences R436: 0.36, R442: 0.29 and R576: 0.81 (Figure 5d), but not in hemimetabolous sequences.

To gain further insight into the differences in conservation across insects, we reduced the multiple sequence alignment dimensionality using a Multiple Correspondence Analysis (MCA), an equivalent of PCA for categorical variables (Lebart, et al. 1984). We performed the dimensionality reduction for the full-length Oskar sequence alignment as well as for the LOTUS and OSK alignments (Supplementary Figure S7). Interestingly, we found that most of the variance in sequence space was due to dipterans and hymenopterans (Supplementary Figure S7). When we considered the OSK domain only, we identified clusters of *Drosophilidae*, *Culicidae* and *Formicidae* sequences (Supplementary Figure S7). This clustering is also reflected for the LOTUS domain, where the *Drosophilidae* and *Culicidae* contribute to a high amount of variance in the first MCA dimension. However, for the LOTUS domain, the *Formicidae* sequences do not cluster away from other Oskar sequences (Supplementary Figure S7). This suggests that the LOTUS domain of Diptera diverged in sequence between *Drosophilidae* and *Culicidae*.

### Evidence for evolution of stronger dimerization potential of the Oskar LOTUS domain in Holometabola

The LOTUS domain dimerizes *in vitro* through electrostatic and hydrophobic contacts of Arg215 of the ꞵ2 sheet and Thr195, Asp197 and Leu200 of the α2 helix (Jeske, et al. 2015; Yang, et al. 2015). To date, however, the biological significance of Oskar dimerization remains unknown. Moreover, the dimerization of the LOTUS domain does not appear to be conserved across all Oskar sequences (Jeske, et al. 2015). Specifically, ten LOTUS domains from non-drosophilid species were tested for dimerization, and only LOTUS domains from *Drosophilidae*, *Tephritidae* and *Pteromalidae* formed homodimers (Jeske, et al. 2015). The other sequences tested, from *Culicidae*, *Formicidae* and *Gryllidae*, remained monomeric under the tested conditions (Jeske, et al. 2015). We selected the LOTUS sequences in our alignment from those six families and placed them into one of two groups, dimeric and monomeric LOTUS, under the assumption that any sequence from that family would conserve the dimerization (or absence thereof) properties previously reported (Jeske, et al. 2015). We asked whether we could detect any evolutionary changes between the two groups in properties of known important dimerization interfaces and residues in our sequence alignment (Jeske, et al. 2015).

In the *D. melanogaster* structure, two key amino acids, D197 and R215, are predicted to form hydrogen bonds that stabilize the dimer (Jeske, et al. 2015). We found that in the dimer group, the electrostatic properties of these two amino acids are highly conserved (-0.75 for D197 and 0.81 for R215), while in the monomer group the electrostatic interaction is not conserved (0.03 for D197 and -0.11 for R215) (Figure 6e). Given the differential conservation between the two groups, our results support the previous finding that disrupting this interaction prevents dimerization (Jeske, et al. 2015). L200 was previously hypothesized to stabilize the interface via hydrophobic forces (Jeske, et al. 2015). We observed that the hydrophobicity of this residue is highly conserved in the dimer group (L200: 0.89), but that in the monomer group this residue is hydrophilic (L200: 2.33) (Figure 6f). In sum, our analyses show that key amino acids in the LOTUS domain evolved differently in distinct insect lineages, in a way that may explain why some insect LOTUS domains dimerize and some do not.

**Figure 6:**
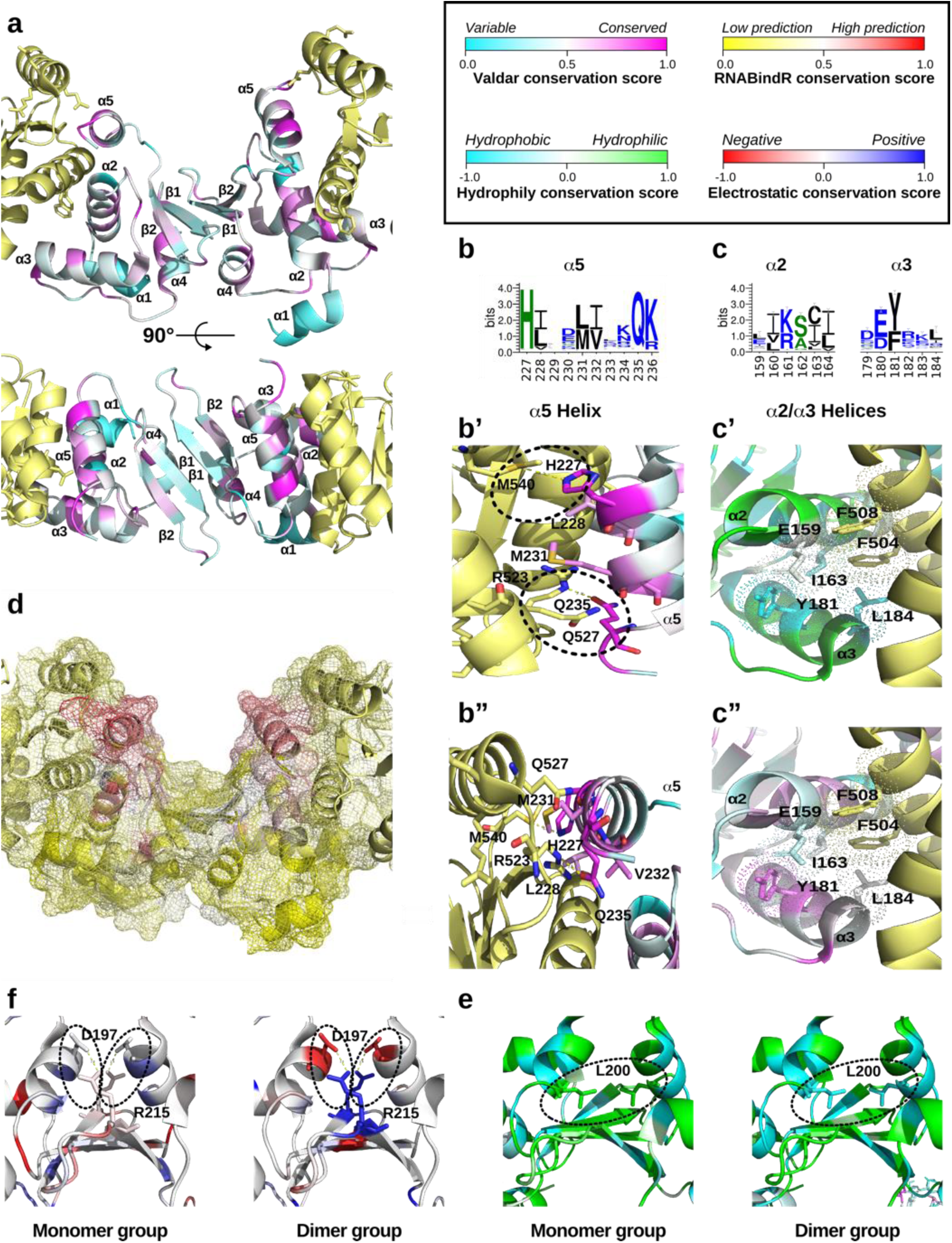
Conservation analysis of the LOTUS domain. **(a)** Ribbon diagram of a LOTUS domain dimer (cyan/magenta) in complex with two Vasa molecules (yellow) (PDBID: 5NT7) from two different angles. Each LOTUS amino acid is colored based on its Valdar conservation score. **(b, c)** Sequence Logo of the α5 and α2/α3 helices respectively generated with WebLogo (Crooks, et al. 2004). Black: hydrophobic residues; blue: charged residues; green: polar residues. **(b’, b”)** Ribbon diagram of the conserved α5 helix, with key amino acids displayed as sticks and colored by Valdar conservation score. Two potential novel Vasa-LOTUS contacts (H227 and Q235) are highlighted with dashed lines. **(c’)** Ribbon diagram of the conserved α2 helix, with key amino acids displayed as sticks and colored by hydrophobicity/hydrophily conservation score. **(c”)** Ribbon diagram of the conserved α2 helix, with key amino acids displayed as sticks and colored by Valdar conservation score. **(d)** Surface mesh rendering colored with the RNABindR RNA binding conservation score. **(e, f)** Ribbon diagram of the LOTUS ꞵ sheet dimerization interface. Left: conservation of monomeric LOTUS domains; right: dimeric LOTUS domains. **(e)** Amino acids colored by electrostatic conservation score. Dashed lines indicate the key electrostatic interaction thought to stabilize the dimerization. **(f)** Amino acids colored by hydrophobicity/hydrophily conservation score. Dashed lines indicate the key hydrophobic pocket thought to stabilize the dimerization.

### Conservation of the Oskar-Vasa interaction interface

Next, we asked whether we could detect differential conservation of the LOTUS-Vasa interface. It was previously reported that the LOTUS domain of Oskar acts as an interaction domain with Vasa (Jeske, et al. 2017), a key protein with a conserved role in the establishment of the animal germ line (Hay, et al. 1990; Lasko 2013). The interaction between Oskar’s LOTUS domain and Vasa is through an interaction surface situated in the pocket formed by the helices α2 and α5 of the LOTUS domain (Figure 6a b and c). Due to the essential role that *vasa* plays in germ line determination (reviewed in Raz 2000; Noce, et al. 2001; Extavour and Akam 2003; Ewen-Campen, et al. 2010; Lasko 2013), and the potential co-option of *oskar* to the germ line determination mechanism in Holometabola (Ewen-Campen, et al. 2012), we hypothesized that evolutionary changes in the conservation of the residues of this interface might be detectable between Holometabola and Hemimetabola. First, we observed that the residues of the LOTUS domain α2 and α5 helices, which directly contact Vasa (Jeske, et al. 2017) were highly conserved overall (α2 average Valdar score 0.49; α5 Valdar score 0.56) (Figure 6b). Specifically, we observed that the previously reported Vasa interacting amino acids A162 and L228 of the LOTUS domain were highly conserved (Valdar score: 0.64 for both residues) (Jeske, et al. 2017). We also noted that Q235 and H227 of the LOTUS domain α5 helix are likely to be important interaction partners due to their high conservation (Valdar score: 0.90 and 0.90 for both residues) (Figure 6b). Moreover, facing the LOTUS domain H227 is Vasa M540, which may act as a proton donor to form a hydrogen bond between the histidine ring and the sulfur atom of the methionine (Pal and Chakrabarti 2001) (Figure 6b and b’). The LOTUS domain α2 helix is overall slightly less conserved than the LOTUS domain α5 helix (Valdar score: 0.49 vs 0.56) (Figure 6a, b’’, c’’), but hydrophobic properties are conserved on one side of the α2 helix (Figure 6c, c’) forming a motif of conserved amino acid properties (Figure 6c’’).

Previous reports have hypothesized that the *D. melanogaster* LOTUS domain could act as a dsRNA binding domain (Anantharaman, et al. 2010; Callebaut and Mornon 2010). However, in *D. melanogaster*, it was later reported that the LOTUS domain did not bind to nucleotides (Jeske, et al. 2015). Therefore, using our dataset we assessed the potential RNA binding properties of LOTUS domains to test the conservation of this prediction. We used the RNABindR algorithm (Terribilini, et al. 2007) to predict potential RNA binding sites of the LOTUS domain, and computed a conservation score for each position (Terribilini, et al. 2007). We found that the α5 helix is the location in the LOTUS domain that has the most conserved prediction for RNA binding (Figure 6d).

Finally, we asked whether the secondary structure of the LOTUS domain might be conserved. Secondary structures are often indicative of the tertiary structure of a domain. Therefore, we reasoned that the secondary structure might be conserved even if the sequence varies. We submitted the LOTUS sequences from all identified Oskar orthologs to the Jpred4 servers (Drozdetskiy, et al. 2015) for secondary structure prediction and mapped the results onto the Oskar alignment we obtained. We found that the secondary structure of LOTUS is highly conserved throughout Oskar orthologs, with the exception of the α1 helix (Supplementary Figure S8) which displays a low conservation score of 0.19 (Figure 6a).

### The core of the OSK domain is conserved

We asked whether the OSK domain showed any differential conservation across the different parts of the domain. We found that the OSK domain of Oskar showed an overall conservation across all insects, similar to the LOTUS domain (Valdar score: 0.51) (Figure 7a). However, the conservation pattern is higher in the core amino acids (Valdar score average of core amino acid: 0.54) when compared to the residues at the surface (Valdar score average for surface amino acid: 0.23) (Figure 7a). Despite the overall low conservation of the residues at the surface of the OSK domain, we found that the electrostatic properties are conserved overall (electrostatic conservation score >0; conserved) in the previously reported putative RNA binding pocket (Yang, et al. 2015). However, as previously mentioned, this conservation is stronger in holometabolous sequences (Figure 5d). These results are in accordance with the potential role of OSK as an RNA Binding domain in the context of germ plasm assembly (Jeske, et al. 2015; Yang, et al. 2015). We also submitted the OSK sequences to the same secondary structure analysis performed on LOTUS. We found that, as for the LOTUS domain, the secondary structure of OSK is highly conserved throughout all insect sequences analyzed (Supplementary Figure S8).

**Figure 7:**
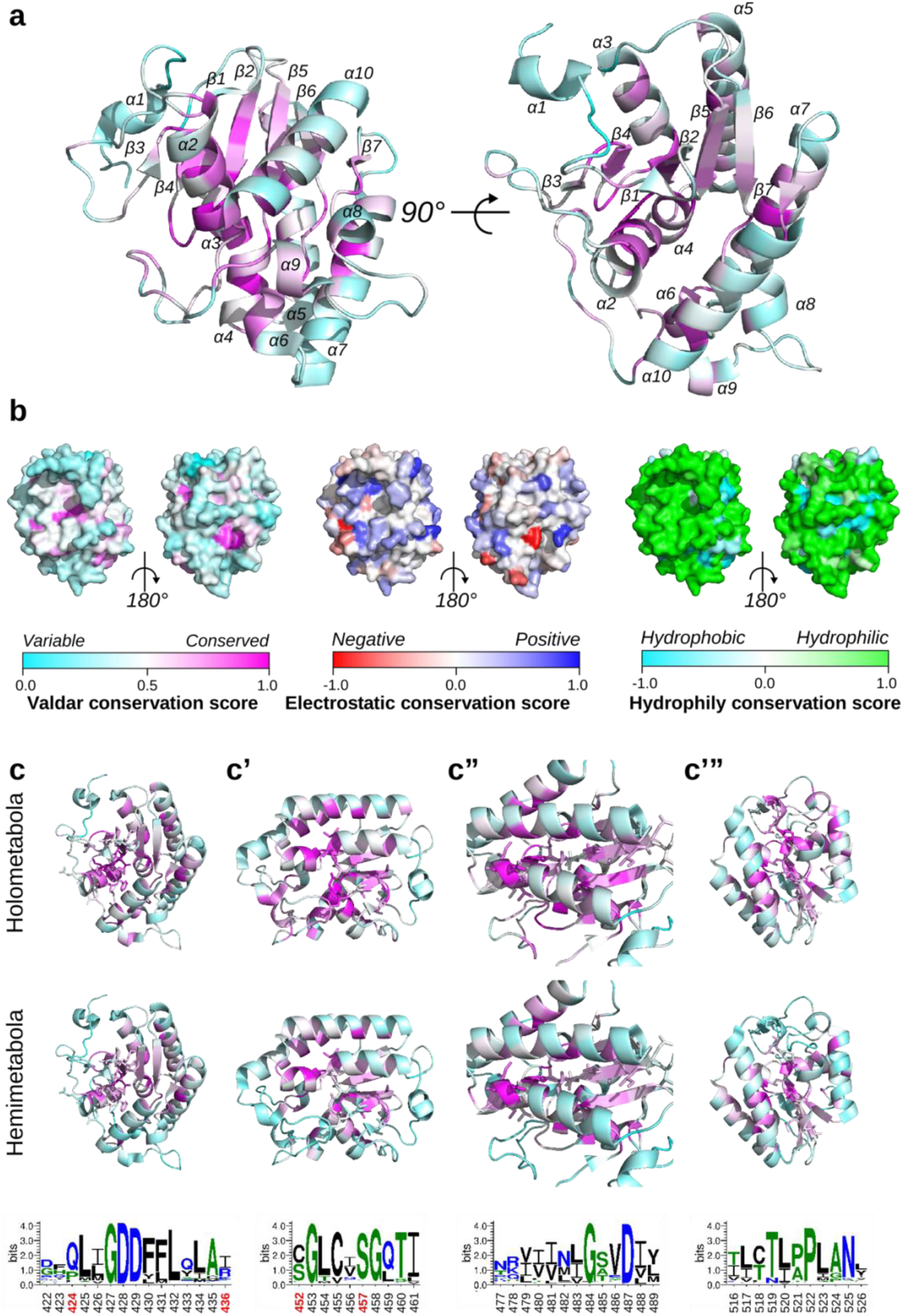
Conservation analysis of the OSK domain. **(a)** Ribbon diagram of the OSK domain (PDBID: 5A4A) from two different angles. Each amino acid is colored based on its Valdar conservation score.**(b)** Protein surface representation of the OSK domain colored by Valdar conservation, electrostatic conservation and hydrophobicity/hydrophily conservation score. **(c, c’, c’’, c’’’)** Ribbon diagram of newly detected conserved motifs of the OSK domain, showing sequence Logo (bottom row) residues as sticks. Each amino acid is colored with Valdar conservation scores of holometabolous (top row) and hemimetabolous (middle row) OSK sequences. Bottom row: sequence Logos of each conserved motif generated with WebLogo (Crooks, et al. 2004). Black: hydrophobic residues; blue: charged residues; green: polar residues. Red numbers: amino acid locations of *D. melanogaster* loss of function *oskar* alleles leading to the loss of *oskar* localization to the posterior pole during embryogenesis (P425S = *osk[8]* (Kim-Ha, et al. 1991); S452L = *osk[255]* = *osk[7]* (Lehmann and Nüsslein-Volhard 1986; Kim-Ha, et al. 1991); S457F = *osk[6B10]* (Breitwieser, et al. 1996)) or to reduced RNA-binding affinity of the OSK domain (R436E (Yang, et al. 2015)).

We then asked if the conservation patterns observed at the core of OSK were clustered in sequence motifs. When we looked at the location of the highly conserved amino acids, we found that the conservation was driven by four well-defined sequence motifs (Figure 7c, c’, c’’, c’’’). Given that *oskar* plays different roles in Holometabola and Hemimetabola, we asked whether the conserved OSK motifs showed any difference in conservation between these two groups. Of the four highly conserved OSK core motifs (Figure 7c, c’, c’’, c’’’), two of them (Figure 7c: Valdar average score: 0.80 and c’’ Valdar average score: 0.71) were conserved across all insects, but the other two showed differential conservation between the holometabolous and hemimetabolous sequences (Figure 7c’: Valdar score average Holometabola: 0.78; Hemimetabola: 0.58 c’’:’ Valdar score average Holometabola: 0.70, Hemimetabola: 0.55). Finally, we noted that only one of the affected OSK domain residues in known loss of function *oskar* alleles affecting posterior patterning in *D. melanogaster*, S457, is conserved across all insects (Valdar score: 0.86). This suggests that the role of the other previously reported important amino acids in the function of *D. melanogaster* OSK (Yang, et al. 2015) might not be conserved in other insects (red positions in Figure 7c, c’, c’’, c’’’).

## Discussion

### An expanded collection of oskar orthologs

*oskar* provides a powerful case study of functional evolution of a gene with an unusual genesis (Blondel, et al. 2020). Here, we gathered the most extensive set of orthologous *oskar* sequences to date. However, most insect genomic and transcriptomic data have been generated from only a few orders, and the vast majority from the Holometabola. Diptera, Lepidoptera, Coleoptera, Hymenoptera and Hemiptera represent 82% of the datasets available at the time of this analysis. We emphasize that expanded taxon sampling, particularly for the Hemimetabola, will be critical for further studies of the evolution of protein function across insects. Moreover, only a small proportion (27% for tissue type, 26% for organism stage, and 14% for sex) of the TSA datasets contained usable metadata regarding the stage and tissue type sampled. Future standardization of the nature and format of transcriptomic metadata would also be a worthwhile endeavor that could increase the efficiency and efficacy of future work.

### Convergent losses and duplications of oskar in insect evolution

A previous report suggested that *oskar* had been lost from the genome of the silk moth *mori* (Lynch, et al. 2011). Our analysis of 232 datasets across 44 of the 126 described lepidopteran families (Kawahara, et al. 2019) strongly suggests that the loss of *oskar* in the Lepidoptera (butterflies and moths) is not unique to the silk moth, but rather occurred early and repeatedly in lepidopteran evolution. The fact that *oskar* is a component of the oosome at the posterior of the oocyte (the wasp germ plasm analog <u>(Quan, et al. 2019)</u>) and required for germ cell formation in the wasp *Nasonia vitripennis* (Lynch, et al. 2011) implies that a common ancestor of Holometabola had already established an *oskar*-dependent inheritance mode of germ line specification. Therefore, the apparent subsequent loss in nearly all Lepidoptera examined of a gene responsible for the establishment of the germ plasm in other Holometabola might seem unexpected. Few studies have directly addressed the molecular mechanisms of germ cell specification in Lepidoptera. In *B. mori* (Bombicidae), *vasa* mRNA (Nakao 1999) and protein (Nakao, et al. 2006), and the transcripts of one of four *nanos* orthologs (*nanos-O*) (Nakao, et al. 2008), have been detected in a region of ventral cortical cytoplasm in pre-blastoderm stage embryos. As putative primordial germ cells form in this location at later stages (Miya 1958), some authors have speculated that a germ plasm, located ventrally rather than posteriorly, may specify germ cells in this moth (Toshiki, et al. 2000; Nakao, et al. 2008). However, recent knockdown experiments showed that maternal *nanos-O* is dispensable for germ cell formation (Nakao and Takasu 2019), consistent with a zygotic, inductive mechanism. In the butterfly *Pararge argeria* (Nymphalidae), no *oskar* ortholog has been identified in the genome (Carter, et al. 2013), but the transcripts of one of four identified *nanos* orthologs (*nanos-O*) have been detected in a small region of ventral cortical ooplasm, again prompting speculation that this lepidopteran may also deploy a germ plasm (Carter, et al. 2015). We suggest that if these or other Lepidoptera do indeed rely on germ plasm to specify their germ line, they may do so using a germ plasm nucleator other than Oskar. For most studied Lepidoptera, however, classical embryological studies report the first appearance of primordial germ cells at post-blastoderm stages, either from the ventral midline of the cellular blastoderm or early germ band (Woodworth 1889; Tomaya 1902; Sehl 1931; Miya 1953, 1958, 1975; Tanaka 1987), from the coelomic sac mesoderm of the abdomen (Johannsen 1929; Eastham 1930; Saito 1937; Presser and Rutschky 1957; Kobayashi and Ando 1984), or from the primary ectoderm of the caudal germ band (Schwangart 1905; Lautenschlager 1932; Ando and Tanaka 1979; Tanaka 1987; Guelin 1994) (Figure S6). Taken together, these data suggest that an inductive mechanism may operate to specify germ cells in most moths and butterflies. We speculate that the loss of *oskar* from most lepidopteran genomes may have facilitated or necessitated secondary reversion to the hypothesized ancestral inductive mechanism for germ line specification.

Another order with apparent near-total absence of *oskar* orthologs is the Hemiptera (true bugs), whose sister group Thysanoptera (thrips) nevertheless possesses *oskar*. This secondary loss of *oskar* from a last common hemipteran ancestor correlates with the reported post-blastoderm appearance of primordial germ cells in the embryo. Classical studies on most hemipteran species describe germ cell formation as occurring after cellular blastoderm formation, on the inner (yolk-facing) side of the posterior blastoderm surface (Metschnikoff 1866; Witlaczil 1884; Will 1888; Mellanby 1935; Butt 1949; Kelly and Huebner 1989; Heming and Huebner 1994). A notable exception to this is the parthenogenetic pea aphid *Acyrthosiphon pisum*, for which strong gene expression and morphological evidence supports a germ plasm-driven germ cell specification mechanism in both sexual and asexual modes (Miura, et al. 2003; Chang, et al. 2006; Lin, et al. 2014). In contrast, studies of the aphids *Aphis plantoides, A. rosea* and *A. pelargonii* describe no germ plasm, and post-blastoderm germ cell formation (Metschnikoff 1866; Witlaczil 1884; Will 1888). However, the genomes of all aphids studied here, including *A. pisum* and three *Aphis* species, appear to lack *oskar*. This suggests that germ plasm assembly in *A. pisum* either does not require a nucleator molecule or uses a novel non-Oskar nucleator.

In the Hymenoptera (ants, bees, wasps and sawflies), our results strongly suggest that *oskar* was lost from the genome of the last common ancestor of bees and spheroid wasps (Supplementary Figure S9). Our analysis further suggests multiple additional independent losses in as many as 25 other hymenopteran lineages, including some for which good quality RefSeq genomes were available (e.g. the slender twig ant *Pseudomyrmex gracilis* or the wheat stem sawfly *Cephus cinctus* (Supplementary Figure S9). However, it would be premature to draw strong conclusions about the number of independent losses given the predominance of transcriptome data in the Hymenoptera.

In addition to convergent losses of *oskar*, we also found evidence for clade-specific duplications of *oskar* in the Hymenoptera. Seven of the nine families containing these putative duplications are families of parasitoid wasps; the remaining two families are ants (Formicidae) and the group of yellowjackets, hornets, and paper wasps (Vespidae) (Figure 4). The phylogenetic relationships of these groups make it highly unlikely that a duplication occurred only once in their last common ancestor, which would be the last common ancestor of all wasps, bees and ants (i.e. Apocrita, all hymenopterans except sawflies) (Supplementary Figure S9). We suggest that the most parsimonious hypothesis is one of three to five independent duplications of *oskar*, followed by at least nine to 14 independent reversions to a single copy, or total loss of the locus (Supplementary Figure 9).

No notable life history characteristics appear to unite those species with multiple *oskar* orthologs: they include eusocial and solitary, sting-bearing and stingless, parasitoid and non-parasitic insects. To our knowledge, neither is there anything unique about the germ line specification process in Hymenoptera with one or more than one *oskar* ortholog. Most Hymenoptera appear to use a germ plasm-driven mechanism to specify germ cells in early blastoderm stage embryos (Supplementary Figure S9 and references therein), and we identified *oskar* orthologs for all such species described in the embryological literature (Supplementary Figure S9). In the notable example of the honeybee *Apis mellifera*, in which cytological and molecular evidence suggests germ cell arise from abdominal mesoderm (Bütschli 1870; Nelson 1915; Fleig and Sander 1985, 1986; Zissler 1992; Gutzeit, et al. 1993; Dearden 2006), we identified no *oskar* ortholog in its well-annotated genome (Supplementary Figure S9), as noted previously by other authors (Lynch, et al. 2011). However, no major differences in germ plasm or pole cell formation have been reported in species or families of ants or wasps with duplicated *oskar* loci, compared with close relatives that possess *oskar* in single copy (e.g. compare the ants *Solenopsis invicta* (at least 2 *oskars*) and *Aphaenogaster rudis* (1 *oskar*) (Khila and Abouheif 2008), or the pteromalid wasps *Nasonia vitripennis* (1 *oskar*) (Lynch and Desplan 2010; Lynch, et al. 2011; Quan, et al. 2019) and *Otitesella tsamvi* (2 *oskars*). Thus, future studies that independently abrogate the functions of each paralog individually, will be needed to determine the biological significance, if any, of these *oskar* duplications.

### Functional implications of differential conservation of the LOTUS and OSK domains

We have identified novel conserved amino acid positions that we hypothesize are important for the Vasa binding properties of the LOTUS domain and the RNA properties binding of the OSK domain (Figure 6 and 7). Our observation of the conservation of the LOTUS domain α2 helix is consistent with its previously reported importance LOTUS-Vasa binding (Jeske, Müller, and Ephrussi 2017). In the α2 helix, we also observed high conservation of H227 and Q235. The positions of these residues suggest they may contribute to the interaction between Vasa and LOTUS. We suggest they should therefore be the target of future mutational studies.

We also uncovered an interesting new conservation pattern within the OSK domain. The conserved amino acids were more abundant in the core of the domain than on the surface. This differential conservation might be relevant to the acquisition of a germ plasm nucleator role of *oskar* in the Holometabla (Figure 5). We noted that the basic properties of surface residues previously reported for *D. melanogaster* (Yang, et al. 2015) are conserved across insects, which might indicate that the RNA binding properties of OSK observed in *D. melanogaster* (Jeske, et al. 2015; Yang, et al. 2015) are also conserved throughout holometabolous insects. We speculate that the comparatively low amino acid conservation of the surface residues in Holometabolous OSK domains, which nevertheless display highly conserved basic properties, could have allowed greater flexibility in the co-evolution of specific RNA binding partners for the OSK domains of different lineages.

### OSK evolved differentially between holometabolous and hemimetabolous insects

Finally, we observed a differential conservation of the OSK domain between hemimetabolous and holometabolous insects. Specifically, we found that the OSK sequence was less conserved across the Holometabola than across the Hemimetabola. This observation raises two potential hypotheses regarding the role of the OSK domain in the functional evolution of Oskar. First, perhaps the apparently relaxed purifying selection experienced by OSK in the Holometabola was necessary for the co-option of *oskar* to a germ plasm nucleation role. Second, Oskar might have a function in the hemimetabolous insects that requires strong conservation of OSK. More studies on the roles and biochemical properties of OSK in hemimetabolous insects will be required to test these hypotheses and further our understanding of the biological relevance of this differential conservation.

In conclusion, analysis of the large dataset of novel Oskar sequences presented here provides multiple new testable hypotheses concerning the molecular mechanisms and functional evolution of *oskar*, that will inform future studies on the contribution of this unusual gene to the evolution of animal germ cell specification.

## Materials and Methods

### Lead contact and materials availability

This study did not generate new unique reagents. This study generated new python3 code and supplementary files referred to below, all of which are available https://github.com/extavourlab/Oskar_Evolution. Requests for further information and requests for resources and reagents should be directed to and will be fulfilled by Cassandra G. Extavour.

### Experimental model and subject details

This study used no animal model, nor any cell culture lines. However, it used previously generated genomic and transcriptomic datasets. All the information regarding how those datasets were generated can be found on their respective NCBI pages. The list of all the datasets used in this study can be found in the following files: ***genome_insect_database.csv****, **transcriptome_insect_database.csv**, **genome_crustacean_database.csv**,* and ***transcriptome_crustacean_database.csv***.

### Genome and transcriptome preprocessing

We collected all available genome and transcriptome datasets from the NCBI repository registered in September 2019 (Figure 2). NCBI maintains two tiers of genomic data: RefSeq, which contains curated and annotated genomes, and GenBank, which contains non-annotated assembled genomic sequences. Transcriptomes are stored in the Transcriptome Shotgun Assembly (TSA) database, with metadata including details on their origin. Among the registered datasets, five genomes were not yet available, and 40 transcriptomes were only available in the NCBI Trace repository. As they did not comply with the TSA database standards, they were excluded from the analysis. To search for *oskar* orthologs in datasets retrieved from GenBank, we needed to generate *in silico* gene model predictions. We used the genome annotation tool Augustus (Stanke et al. 2006), which requires a Hidden Markov Model (HMM) gene model. To use HMMs producing gene models that would be as accurate as possible for non-annotated genomes, we selected the most closely related species (species with the most recent last common ancestor) that possessed an annotated RefSeq genome. We then used the Augustus training tool to build an HMM gene model for each genome.

We automated this process by creating a series of python scripts that performed the following tasks:

(1) ***1.1_insect_database_builder.py***: This script collects the NCBI metadata regarding genomes and transcriptomes. Using the NCBI Entrez API, it collects the most up to date information on RefSeq, GenBank, and TSA to generate two CSV files: *genome_insect_database.csv* and *transcriptome_insect_database.csv*.
(2) ***1.2_data_downloader.py***: This is a python wrapper around the *rsync* tool that downloads the sequence datasets present in the tables created by (1). It automatically downloads all the available information into a local folder.
(3) ***1.3_run_augustus_training.py***: This is a python wrapper around the Augustus training tool. It uses the metadata gathered using (1) and the sequence information gathered using (2) to build HMM gene models of all RefSeq datasets. It outputs sbatch scripts that can be run either locally, or on a SLURM-managed cluster. Those scripts will create unique HMM gene models per species.

At the time of this analysis (September 2019), 133 insect genomes were collected from the RefSeq database, 309 genomes from the GenBank database, and 1123 transcriptomes from the TSA database. All the accession numbers and metadata are available in the two tables (***genome_insect_database.csv*** and ***transcriptome_insect_database.csv***) provided in the supplementary files. This pipeline was repeated for crustaceans and the information can be found in the following two files: ***genome_crustacean_database.csv*** and ***transcriptome_crustacean_database.csv***.

### Creation of protein sequence databases

The classical approach for orthology detection compares protein sequences to amino acid HMM corresponding to the gene of interest. Since we used three different NCBI databases, we performed the following preprocessing actions:

1) RefSeq: well-annotated genomes from NCBI contain gene model translation; no extra processing was required.
2) GenBank: Using the HMMs created from the RefSeq databases, we created gene models for each GenBank genome using Augustus and a custom HMM gene model. To choose which HMM gene model to use, we selected the one for each insect order that had the highest training accuracy. In the case where an insect order did not have any member in the RefSeq database, we used the model of the most closely related order. We then translated the inferred coding sequences to create a protein database for each genome. The assignment of the models used to infer the proteins of each GenBank genome is available in the ***Table_S4_models.csv*** available through the GitHub repository for this study at https://github.com/extavourlab/Oskar_Evolution. To automate the process, we created a custom python script available in the file **1.4_run_augustus.py**.
3) TSA: Transcriptomes were translated using the emboss tool Transeq (Madeira, et al. 2019). We used this tool with the default parameters, except for the six-frame translation, trim and clean flags. This generated amino acid sequences for each transcript and each potential reading frame.

### Identification of oskar orthologs

The *oskar* gene is composed of two conserved domains, LOTUS and OSK, separated by a highly variable interdomain linker sequence (Ahuja and Extavour 2014; Jeske, et al. 2015; Yang, et al. 2015). To our knowledge, no other gene reported in any domain of life possesses this domain composition (Blondel, et al. 2020). Therefore, here we use the same definition of *oskar* orthology as in our previous work: a sequence possessing a LOTUS domain followed by an interdomain region, and then an OSK domain (Blondel, et al. 2020). To maximize the number of potential orthologs, we searched each sequence with the previously generated HMM for the LOTUS and OSK domains (Blondel, et al. 2020). The presence and order of each domain were then verified for each potential hit and only sequences with the previously defined Oskar structure were kept for further processing. We used the HMMER 3.1 tool suite to build the domain HMM (*hmmbuild* with default parameters), and then searched the generated protein databases (see *Creation of protein sequence databases* above) using those models (*hmmsearch* with default parameters). Hits with an E-value >= 0.05 were discarded. A summary of all searches performed is compiled in ***Table_S5_searches.csv*** in the GitHub repository for this study at https://github.com/extavourlab/Oskar_Evolution.

All the hits were then aligned with *hmmalign* with default parameters and the HMM of the full-length Oskar alignment previously generated (Blondel, et al. 2020). The resulting sequences were automatically processed to remove assembly artifacts, and potential isoforms. This filtration step was automated and went as follows: First, the sequences were grouped by taxon. Then each group of sequences was aligned using MUSCLE (Edgar 2004) with default parameters. The Hamming distance (Hamming 1950), a metric that computes the number of different letters between two strings, between each sequence in the alignment, was computed. If any group of sequences had a Hamming distance of >80%, then we only kept the sequence with the lowest E-value match. This created a set of sequences containing multiple *oskar* orthologs per species only if they were the likely product of a gene duplication event. We then used the resulting new alignment to generate a new domain HMM and a new full-length Oskar HMM (using *hmmbuild* with default parameters) and ran further iterations of this detection pipeline until we could detect no new *oskar* orthologs in the available sequence datasets. We called this final set the **filtered set** of sequences and used it in all subsequent orthology analyses unless otherwise specified.

The Oskar sequences obtained are available in the following supplementary files: ***Oskar_filtered.aligned.fasta*, *Oskar_filtered.fasta* and *Oskar_consensus.hmm*.** The domain definitions for the LOTUS and OSK domains are available in the following supplementary files: ***Oskar_filtered.aligned.LOTUS_domain.fasta*, *LOTUS_consensus.hmm*, *Oskar_filtered.aligned.OSK_domain.fasta*, *OSK_consensus.hmm* (see *1.5_Oskar_tracker.ipynb*)**.

### Correlative analysis of assembly quality and absence of oskar

Using the metadata gathered previously from NCBI databases (see *Genomes and transcriptomes preprocessing* above) we created two pools of source data: genomes where we identified an *oskar* sequence, and genomes where we failed to find a sequence that met our orthology criteria. We then compared the two distributions for each of the 8 available assembly statistics: (1) Contig and (2) Scaffold N50, (3) Contig and (4) Scaffold L50, (5) Contig and (6) Scaffold counts, and (7) Number of Contigs and (8) Scaffolds per genome length. Finally, we performed a Mann-Whitney U statistical analysis to compare the means of the two distributions (see ***2.1_Oskar_discovery_quality.ipynb***).

### TSA metadata parsing and curation

Datasets in the TSA database are associated with a biosample object that contains all the metadata surrounding the RNA sequencing acquisitions. These metadata can include information about one or both the tissue of origin and the organism’s developmental stage. We first automated the retrieval of these metadata using a custom python script that used the NCBI Entrez API (see ***2.3_Oskar_tissues_stages.ipynb***). However, the metadata proved to be complex to parse for the following reasons: (1) not all projects had the data entered in the corresponding tag, (2) some data contained typographical errors, and (3) multiple synonyms were used to describe the same thing with different words in different datasets. We therefore created a custom parsing and cleaning pipeline that corrected mistakes and aggregated them into a cohesive set of unique terms that we thought would be most informative to interpret the presence or absence of *oskar* orthologs (see ***2.3_Oskar_tissues_stages.ipynb*** to see the mapping table). This strategy sacrificed some of the fine-grained information contained in custom metadata (for example “right leg” became “leg”) but allowed us to analyze the expression of *oskar* using consistent criteria throughout all the datasets. This pipeline generated, for all available datasets, a table of tissues and developmental stages including *oskar* presence or absence in the dataset (see ***Oskar_all_tissues_stages.csv***).

### Dimensionality reduction of Oskar alignment sequence space

The Oskar alignment was subjected to a Multiple Correspondence Analysis (MCA). Similar to a PCA, dimension vectors were first computed to maximize the spread of the underlying data in the new dimensions, except that instead of a continuous dataset, each variable (here an amino acid at a given position) contributes to the continuous value on that dimension. Once the projection vectors are computed, each sequence was then mapped onto the dimensions. Each amino acid position (column) in the alignment was considered a dimension with a possible value set of 21 (20 amino acids and gap). We first removed the columns of low information (columns that had less than 30% amino acid occupancy) using trimal (Capella-Gutierrez, et al. 2009) with a cutoff parameter set at 0.3. Then, the alignment was decomposed into its eigenvectors, and projected to the first three components. To perform this decomposition, we implemented a previously developed preprocessing method (Rausell, et al. 2010) in a python script (see ***MCA.py*** and ***2.8_Oskar_MCA_Analysis.ipynb***) and performed the eigenvector decomposition with the previously developed MCA python library (see *Key Resource Table*). We ran the same algorithm on the LOTUS domain, OSK domain, and full-length Oskar alignments obtained above (see *Identification of* oskar *orthologs* above).

### Phylogenetic inference of Oskar sequences in the Hymenoptera

We aligned all hymenopteran Oskar sequences using PRANK (Loytynoja 2014) with default parameters. We then manually annotated duplicated sequences by considering two sequences from the same species that had < 80% amino acid identity, as within-species duplications of *oskar*. We trimmed this alignment to remove all columns with less than 50% occupancy using trimal with the cutoff parameter set at 0.5. To reconstruct the phylogeny of these sequences, we used the maximum likelihood inference software RAxML (Stamatakis 2014) with a gamma-distributed protein model, and activated the flag for auto model selection. We ran 100 bootstraps and then visualized and annotated the obtained tree with Ete3 (Huerta-Cepas, et al. 2016) in a custom ipython notebook (see ***2.7_Oskar_duplication.ipynb***).

### Calculation of Oskar conservation scores

Using the large set of orthologous Oskar sequences obtained as described above, we computed different conservation scores for each amino acid position. This methodology relies on the hypotheses that if an amino acid, or its associated chemical properties at a particular position in the sequence are important for the structure and/or function of the protein, they will be conserved across evolution. We considered multiple conservation metrics, each highlighting a particular aspect of the protein’s properties as described in the followng sections. The scores can be found in the supplementary file ***scores.csv***.

### Computation of the Valdar score

The Valdar score (Valdar 2002) attempts to account for transition probabilities, stereochemical properties, amino acid frequency gaps, and, particularly essential for this study, sequence weighting. Due to the heterogeneity of sequence dataset availability, most Oskar sequences occupy only a small portion of insect diversity, primarily Hymenoptera, and Diptera. Sequence weighting allows for the normalization of the influence of each sequence on the score based on how many similar sequences are present in the alignment (Valdar 2002). We implemented the algorithm described in (Valdar 2002) in a python script (see ***besse_blondel_conservation_scores.py***), then calculated the conservation scores for the Oskar alignment we generated above.

### Computation of the Jensen-Shannon Divergence score

Jensen-Shannon Divergence (JSD) (Lin 1991; Capra and Singh 2007) uses the amino acid and stereochemical properties to infer the “amount” of evolutionary pressure an amino acid position may be subject to. This score uses an information theory approach by measuring how much information (in bits) any position in the alignment brings to the overall alignment (Capra and Singh 2007). This score also takes into account neighboring amino acids in calculating the importance of each amino acid. We used the previously published python code to calculate the JSD of our previously generated Oskar alignment (Capra and Singh 2007) (see ***score_conservation.py***).

### Computation of the Conservation Bias

The measure of differences in conservation between the holometabolous and hemimetabolous Oskar sequences presented in the results was done as follows: we first split the alignment into two groups containing the sequences from each clade (see ***2.4_Oskar_pgc_specification.ipynb***). Due to the high heterogeneity in taxon sampling between hemimetabolous and holometabolous insects, we ran a bootstrapped approximation of the conservation scores on holometabolous sequences. We randomly selected N sequences (N = the number of hemimetabolous sequences), computed the Valdar conservation score (see *Computation of the Valdar score* above), and stored it. After 1000 iterations, we computed the mean conservation score for each position for holometabolous sequences. For hemimetabolous sequences, we directly calculated the Valdar score using the method as described above (see *Computation of the Valdar score* above). For each position, we then computed what we refer to as the “conservation bias” between Holometabola and Hemimetabola by taking the ratio of the log of the conservation score Holometabola and Hemimetabola. Conservation Bias = log(Valdar_holo_) / log(Valdar_hemi_) for each position (see ***3.4_LogRatio_Bootstrap.ipynb***).

### Computation of the electrostatic conservation score

To study the conservation of electrostatic properties of the Oskar protein we computed our own implementation of an electrostatic conservation score (see ***besse_blondel_conservation_scores.py***). Aspartic acid and Glutamic acid were given a score of -1, Arginine and Lysine a score of 1, and Histidine a score of 0.5. All other amino acids were given a score of 0. Then, we summed the electrostatic score for each sequence at each position and divided this raw score by the total number of sequences in the alignment. This computation assigns a score between -1 and 1 at each position, - 1 being a negative charge conserved across all sequences, and 1 a positive charge.

### Computation of the hydrophobic conservation score

To study the conservation of hydrophobic properties of the Oskar protein we implemented our own hydrophobic conservation score (see ***besse_blondel_conservation_scores.py***). At each position, each amino acid was given a hydrophobic score taken from a previously published scoring table (Moon and Fleming 2011). (This table is implemented in the ***besse_blondel_conservation_score.py*** file for simplicity.) Scores at each position were then averaged across all sequences. This metric allowed us to measure the hydrophobicity conservation of each position in the alignment and is bounded between 5.39 and -2.20.

### Computation of the RNA binding affinity score

RNA binding sites are defined as areas with positively charged residues and hydrophobic residues. To estimate the conservation of RNA binding sites in *oskar* orthologs, we used RNABindR v2.0 (Terribilini, et al. 2007), an algorithm predicting putative RNA binding sites based on sequence information only. We automated the calculation for each sequence by writing a python script that submitted a request to the RNABindR web service (see ***RNABindR_run_predictions.py***). We then aggregated all results into a scoring matrix, and averaged the score obtained for each position. We call this score the RNABindR score and hypothesize that it reflects the conservation of RNA binding properties of the protein. Importantly, this score was obtained in 2017 for only a subset of 219 proteins used in this study (indicated in the supplementary files at: 03_Oskar_scores_generation/RNABindR_raw_sources). Since then, the RNABindR server has been defunct and we could not repeat those measurements as the source code for this software is unavailable.

### Computation of secondary structure conservation

Due to the overall low conservation of the LOTUS domain, we decided to see whether the secondary structure was conserved. To this end, we used the secondary structure prediction algorithm JPred 4 (Drozdetskiy, et al. 2015). Given an amino acid sequence, this tool returns a positional prediction for α-helix, β-sheet or unstructured. We used the JPred4 web servers to compute the predictions and processed them into a secondary structure alignment (see ***2.6_Oskar_lotus_osk_structures.ipynb***). We then used WebLogo (Crooks, et al. 2004) to visualize the conservation of the secondary structure.

### Visualization of conservation scores

We used PyMOL (DeLano 2002) to map the computed conservation scores onto the solved structures of LOTUS and OSK (Jeske, et al. 2015; Jeske, et al. 2017). At the time of writing, no full-length Oskar protein structure had been reported. With the caveat that all visualization was done on the structure of the *Drosophila melanogaster* protein domains, we created a custom python script that augments PyMOL with automatic display and coloring capacities. This script is available as ***Oskar_pymol_visualization.py***, and contains a manual at the beginning of the file. For the OSK domain, we used the structure PDBID: 5A4A, and for the LOTUS domain, PDBID: 5NT7 (Jeske, et al. 2015; Jeske, et al. 2017). The LOTUS structure we used is in complex with Vasa, and in a dimeric form (Jeske, et al. 2017), allowing for easy interpretation of the different conservation scores. For the OSK structure, we removed the residues 399-401 and 604-606 from the PDB file as those amino acids did not align across all sequences and therefore showed highly biased conservation scores.

### Statistical analysis

All statistical analyses were performed using the scipy stats module (https://www.scipy.org/). Significance thresholds for p-values were set at 0.05. Statistical tests and p-values are reported in the figure legends. All statistical tests can be found in the ipython notebooks mentioned below.

### Data and code availability

The study generated a series of python 3 script and python 3 ipython notebook files that perform the entire analysis. All the results presented in this paper can be reproduced by running the aforementioned python 3 code. The primary data, *oskar* orthologs, Oskar alignments, trees, and conservation statistics as well as the code created and used are available as supplementary information. For ease of access, legibility, and reproducibility, the code and datasets have been deposited in a GitHub repository available at https://github.com/extavourlab/Oskar_Evolution.

### Software and libraries

All software and libraries used in this study are published under open source libre licenses and are therefore available to any researcher.

| Type | Name | Version | Source |
| --- | --- | --- | --- |
| Software | HMMER | 3.1.b2 | http://hmmer.org/ |
| Software | PyMOL | 1.8.x | https://pymol.org |
| Software | rsync | 3.1.2 | http://rsync.samba.org/ |
| Software | Python 3 | 3.7 | https://www.python.org/ |
| Software | MrBayes | 3.2.6 | http://nbisweden.github.io/MrBayes/ |
| Software | trimal | 1.2rev59 | http://trimal.cgenomics.org/ |
| Software | transeq | 6.6.0.0 | http://emboss.sourceforge.net/apps/cvs/emboss/apps/transeq.html |
| Software | augustus | 2.5.5 | http://augustus.gobics.de/ |
| Software | JPred4 | 4.0 | http://www.compbio.dundee.ac.uk/jpred/ |
| Software | RNABindR | 2.0 | http://ailab1.ist.psu.edu/RNABindR/ |
| Software | Inkscape | 0.92.3 | https://inkscape.org/ |
| Library | jupyter | 4.4.0 | https://jupyter.org/ |
| Library | ete3 | 3.3.1 | http://etetoolkit.org |
| Library | pandas | 0.25.1 | https://pandas.pydata.org/ |
| Library | mca | 1.0.3 | https://pypi.org/project/mca/ |
| Library | fuzzywuzzy | 0.17.0 | https://github.com/seatgeek/fuzzywuzzy |
| Library | BeautifulSoup4 | 4.6.3 | https://pypi.org/project/beautifulsoup4/ |
| Library | biopython | 1.74 | https://pypi.org/project/biopython/ |
| Library | numpy | 1.16.2 | https://www.numpy.org/ |
| Library | seaborn | 0.9.0 | https://seaborn.pydata.org/ |
| Library | matplotlib | 3.0.0 | https://matplotlib.org/ |
| Library | scipy | 1.1.0 | https://www.scipy.org/ |
| Library | progressbar | 3.38.0 | https://github.com/niltonvolpato/python-progressbar |

## Acknowledgements

This work was supported by funds from Harvard University, and support to SB from the Master’s in Bioinformatics Program of the University of Bordeaux. We thank members of the Extavour lab for discussion.

## Supplementary Materials

These Supplementary Materials contain the following:

- Supplementary Tables S1 through S5
- Supplementary Figures S1 through S9

## Supplementary Table Legends

**Supplementary Table S1:** Number of *oskar* sequences identified per order and per data source. Each row corresponds to an order and a data source: GCF: RefSeq; GCA: GenBank, TSA: Transcriptome Shotgun Assembly Database. “Filtered hits” column indicates the number of hits after the filtration algorithm described in the Methods is applied. Rightmost column defines the proportion of *oskar* sequences identified, as the number of datasets with a filtered hit divided by the total number of datasets searched.

| **Insect Order** | **Source** | **Number of datasets searched** | **Total hits** | **Filtered hits** | **% of datasets with oskar identified** |
| --- | --- | --- | --- | --- | --- |
| Archaeognatha | GCA | 1 | 0 | 0 | 0 |
| Archaeognatha | TSA | 2 | 0 | 0 | 0 |
| Blattodea | GCA | 3 | 1 | 1 | 33.33 |
| Blattodea | GCF | 2 | 0 | 0 | 0 |
| Blattodea | TSA | 51 | 7 | 7 | 13.73 |
| Coleoptera | GCA | 12 | 1 | 1 | 8.33 |
| Coleoptera | GCF | 9 | 3 | 2 | 22.22 |
| Coleoptera | TSA | 86 | 31 | 14 | 16.28 |
| Collembola | TSA | 9 | 0 | 0 | 0 |
| Dermaptera | TSA | 7 | 0 | 0 | 0 |
| Diptera | GCA | 115 | 63 | 60 | 52.17 |
| Diptera | GCF | 43 | 58 | 43 | 100 |
| Diptera | TSA | 162 | 72 | 58 | 35.8 |
| Embioptera | TSA | 5 | 0 | 0 | 0 |
| Ephemeroptera | GCA | 2 | 0 | 0 | 0 |
| Ephemeroptera | TSA | 5 | 1 | 1 | 20 |
| Grylloblattodea | TSA | 2 | 0 | 0 | 0 |
| Hemiptera | GCA | 18 | 0 | 0 | 0 |
| Hemiptera | GCF | 12 | 0 | 0 | 0 |
| Hemiptera | TSA | 192 | 1 | 0 | 0 |
| Hymenoptera | GCA | 52 | 32 | 30 | 57.69 |
| Hymenoptera | GCF | 47 | 36 | 32 | 68.09 |
| Hymenoptera | TSA | 301 | 157 | 128 | 42.52 |
| Lepidoptera | GCA | 80 | 0 | 0 | 0 |
| Lepidoptera | GCF | 17 | 0 | 0 | 0 |
| Lepidoptera | TSA | 135 | 24 | 4 | 2.96 |
| Mantodea | TSA | 13 | 0 | 0 | 0 |
| Mantophasmatodea | TSA | 2 | 0 | 0 | 0 |
| Mecoptera | TSA | 4 | 2 | 2 | 50 |
| Megaloptera | TSA | 3 | 0 | 0 | 0 |
| Neuroptera | TSA | 7 | 1 | 1 | 14.29 |
| Odonata | GCA | 2 | 0 | 0 | 0 |
| Odonata | TSA | 7 | 0 | 0 | 0 |
| Orthoptera | GCA | 3 | 0 | 0 | 0 |
| Orthoptera | TSA | 28 | 2 | 2 | 7.14 |
| Phasmatodea | GCA | 13 | 0 | 0 | 0 |
| Phasmatodea | TSA | 31 | 6 | 2 | 6.45 |
| Phthiraptera | GCF | 1 | 0 | 0 | 0 |
| Phthiraptera | TSA | 7 | 0 | 0 | 0 |
| Plecoptera | GCA | 3 | 0 | 0 | 0 |
| Plecoptera | TSA | 8 | 3 | 3 | 37.5 |
| Psocoptera | TSA | 23 | 5 | 5 | 21.74 |
| Raphidioptera | TSA | 3 | 0 | 0 | 0 |
| Siphonaptera | GCF | 1 | 0 | 0 | 0 |
| Siphonaptera | TSA | 4 | 0 | 0 | 0 |
| Strepsiptera | GCA | 1 | 0 | 0 | 0 |
| Strepsiptera | TSA | 2 | 0 | 0 | 0 |
| Thysanoptera | GCA | 1 | 1 | 1 | 100 |
| Thysanoptera | GCF | 1 | 1 | 1 | 100 |
| Thysanoptera | TSA | 11 | 10 | 8 | 72.73 |
| Trichoptera | GCA | 3 | 1 | 1 | 33.33 |
| Trichoptera | TSA | 7 | 2 | 2 | 28.57 |
| Zoraptera | TSA | 2 | 0 | 0 | 0 |
| Zygentoma | TSA | 4 | 1 | 1 | 25 |
| Crustacea | TSA | 168 | 0 | 0 | 0 |
| Crustacea | GCF | 1 | 0 | 0 | 0 |
| Crustacea | GCA | 11 | 0 | 0 | 0 |

**Supplementary Table S2:** Genome quality correlation to *oskar* identification. Mean and median values for the distributions of each indicated genome quality parameter, in which *oskar* was (a) or was not (b) identified. The means of both distributions are significantly different for all metrics (Mann Whitney U test, p<0.05). See Supplementary Figure S2 of graphical representation of distributions.

|  | **Genome parameter** | **(a) *oskar* Identified** | **(b) *oskar* not identified** | **ratio (a):(b)** |
| --- | --- | --- | --- | --- |
| # contigs | mean | 255,015 | 43,280 | 5.89 |
|  | median | 69,255 | 20,653 | 3.35 |
| # scaffolds | mean | 182,706 | 23,596 | 7.74 |
|  | median | 40,960 | 9,398 | 4.36 |
| contig N50 (bp) | mean | 324,036 | 726,696 | 0.45 |
|  | median | 14,052 | 40,079 | 0.35 |
| scaffold N50 | mean | 2,636,825 | 5,695,299 | 0.46 |
|  | median | 96,730 | 385,460 | 0.25 |
| contig L50 | mean | 40,955 | 3,701 | 11.07 |
|  | median | 6,868 | 1,300 | 5.28 |
| scaffold L50 | mean | 27,269 | 1,500 | 18.18 |
|  | median | 1,131 | 191 | 59.53 |
| # contigs per genome length | mean | 0.00060 | 0.00017 | 3.53 |
|  | median | 0.00021 | 0.00009 | 2.33 |
| # scaffolds per genome length | mean | 0.00045 | 0.00009 | 5.00 |
|  | median | 0.00013 | 0.00005 | 2.60 |

**Supplementary Table S3. Assignment of metadata to germ line or brain categories.** This table is found in ***Data>02_oskar_analyses/2/3/TableS3_germline_brain_table.csv*** at the GitHub repository https://github.com/extavourlab/Oskar_Evolution.

**Supplementary Table S4. Models used to create protein sequence databases.** This table shows which models were used to run the *ab initio* gene detection algorithm Augustus as described in Methods and Materials. Column order corresponds to any GCA dataset of an organism from this order. “Family” column is only used if a member of this order but of a different family was used. Finally, “augustus_model” shows which GCF dataset or premade augustus model, was used to run the gene prediction. This table is found in ***Data>Tables>TableS4_models.csv*** at the GitHub repository https://github.com/extavourlab/Oskar_Evolution.

**Supplementary Table S5. *oskar* search results master table.** This table summarizes all results of the *oskar* search performed on each dataset. Each row corresponds to a dataset. Columns are as follows: Id: the dataset NCBI identifier; Species: the organism’s species name; Family_name: the organism’s family name; Order_name: the organism’s order name; Hits: the number of sequences in the dataset found that satisfy our criteria for *oskar* orthology; Source: the NCBI database from which this dataset was downloaded; Filtered_hits: the number of *oskar* sequins in remaining the dataset after the filtration process was applied to all identified *oskar* sequences. For more information on the criteria used for *oskar* orthology and the filtration process, please see the Materials and Methods “*Identification of oskar orthologs”*. This table is found in ***Data>Tables>TableS5_models.csv*** at the GitHub repository https://github.com/extavourlab/Oskar_Evolution.

## Supplementary Figure Legends

**Supplementary Figure S1:**
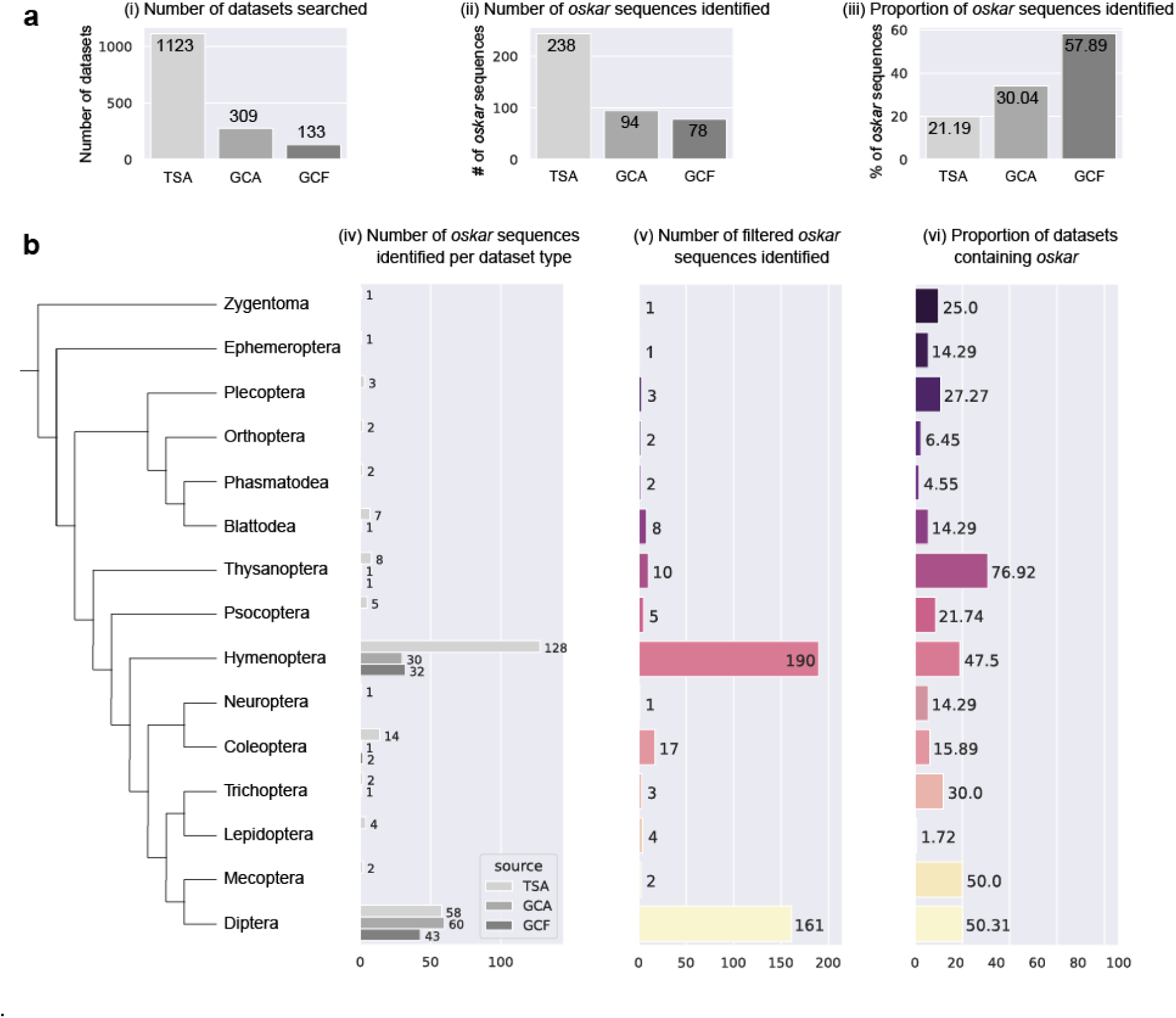
Summary statistics of the search for *oskar* orthologs. **(a)** Summary of searches and results for each of the three sources of data searched, from left to right: (i) The total number of datasets searched from all three sources (TSA: Transcriptome Shotgun Assembly Database; GCA: GenBank; GCF: RefSeq); (ii) the number of filtered *oskar* sequences identified in each of those datasets; and (iii) the proportion of filtered *oskar* sequences identified in each of the three sources. **(b)** Summary statistics broken down by insect orders. Only orders where an *oskar* sequence was identified are shown. From left to right: (iv) The number of *oskar* sequences identified in each of the three data sources; (v) the total number of filtered *oskar* sequences identified per order; (vi) the proportion of all searched datasets per order where an *oskar* sequences was identified. See also Supplementary Table 1

**Supplementary Figure S2:**
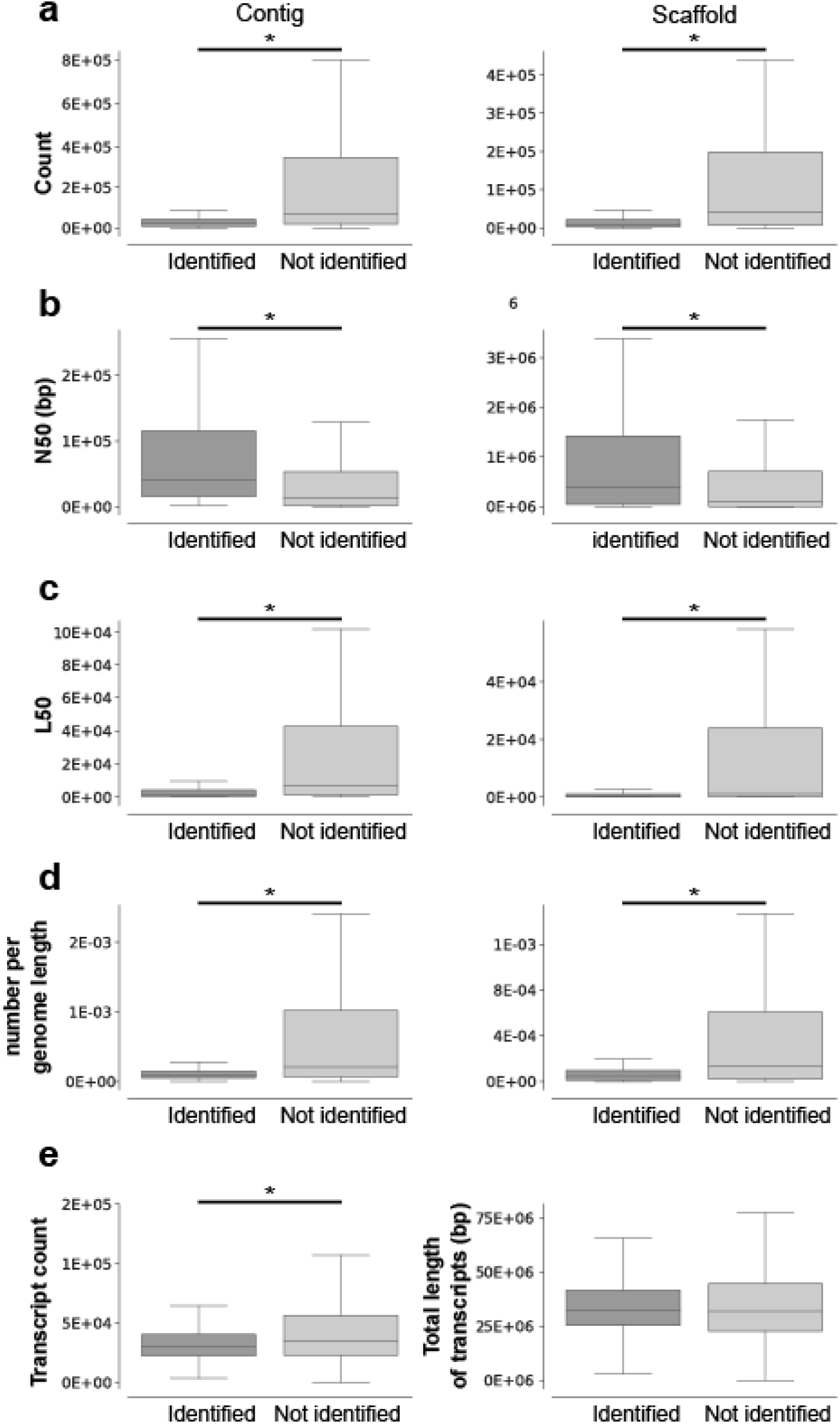
Genome and transcriptome quality correlation to *oskar* identification. Shown are box plots of the distribution of *oskar* orthologs identified (ortholog identified or not identified) with respect to multiple genome and transcriptome quality metrics. For each metric, the means of both distributions were tested for significant differences using a Mann Whitney U test. A bar with an * is displayed if the p-value was less than 0.05. Mean and median values presented in Supplementary Table S2.

**Supplementary Figure S3:**
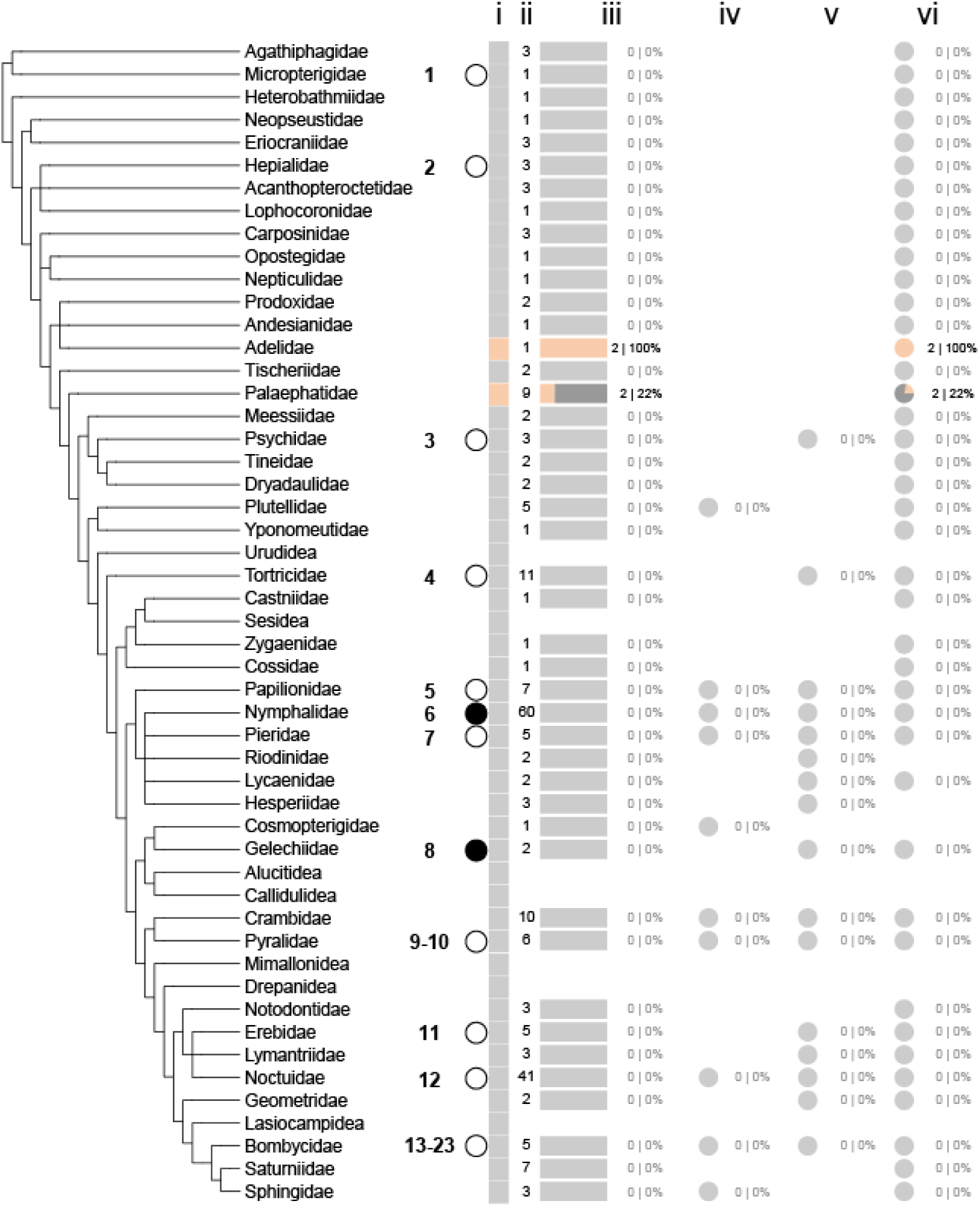
Evidence for loss of *oskar* in Lepidoptera. Phylogeny of the Lepidoptera as per (Kawahara, et al. 2019). Next to each lepidopteran family are shown summary data regarding the status of *oskar* identification in our searches. Symbols with column labels in order from left to right: (i) vertical rectangles: grey: no *oskar* ortholog was identified in this family; range: at least one *oskar* ortholog was identified in this order. (ii) number of datasets searched. (iii) horizontal rectangles: proportion of searched datasets in which an *oskar* ortholog was identified; colors as in (i); numbers and proportions at right. (iv) pie chart: proportion of *oskar* sequences identified in RefSeq (GCF) datasets; numbers and proportions at right. (v) pie chart: proportion of *oskar* sequences identified in GenBank (GCA) datasets; numbers and proportions at right. (vi) pie chart: proportion of *oskar* sequences identified in Transcriptome Shotgun Assembly Database (TSA) datasets; numbers and proportions at right. Circles to the right of some family names indicate that there is literature evidence for involvement of germ plasm (black) or no germ plasm (white) in germ cell specification. Numbers to the left of the circles indicate references to the primary literature as follows: [1]: (Kobayashi and Ando 1984); [2] (Ando and Tanaka 1979); [3] (Lautenschlager 1932); [4] (Anderson and Wood 1968); [5] (Tanaka 1987); [6] (Woodworth 1889); [7] (Eastham 1930); [8] (Berg and Gassner 1978); [9-10] (Sehl 1931; Guelin 1994); [11] (Johannsen 1929); [12] (Presser and Rutschky 1957); [13-23]: (Tomaya 1902; Schwangart 1905; Saito 1937; Miya 1953, 1958, 1975; Nakao 1999; Toshiki, et al. 2000; Nakao, et al. 2006; Nakao, et al. 2008; Nakao and Takasu 2019). No datasets were available for Urudidea, Sesidea, Alucitidea, Callidulidea, Mimallonidea, Drepanidea or Lasiocampidea at the time of analysis.

**Supplementary Figure S4:**
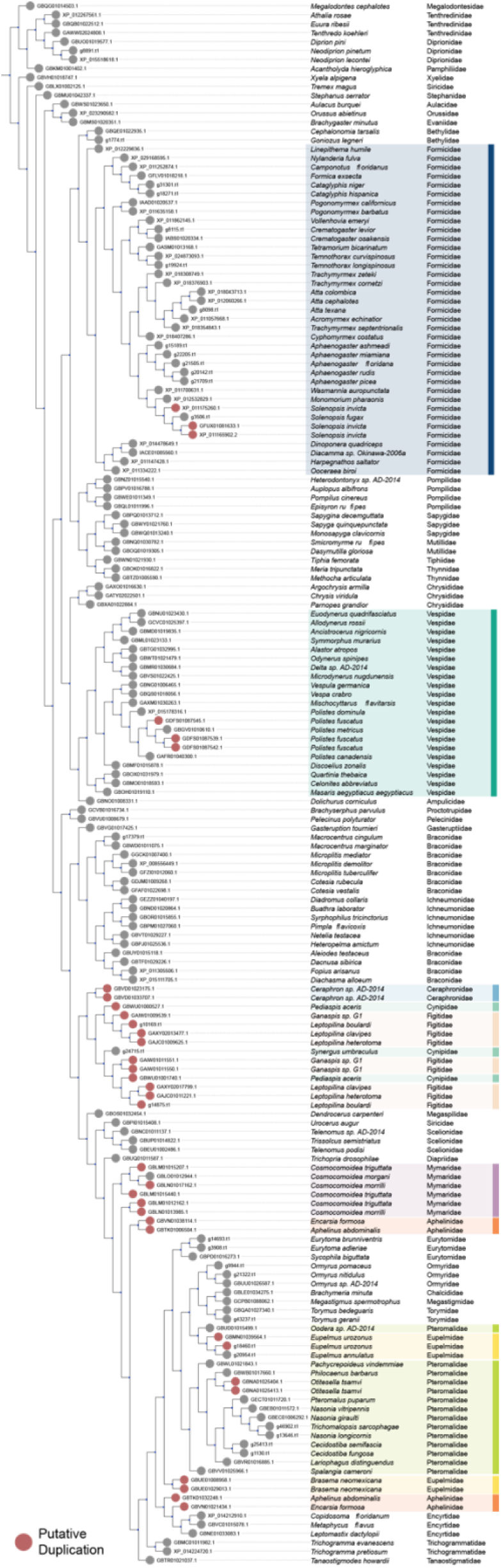
Evidence for duplication of *oskar* in Hymenoptera. Phylogenetic tree of all hymenopteran Oskar sequences inferred using RaxML with 100 bootstraps. Branch length normalized to show only the topology. Each leaf is an Oskar ortholog. Gray: only one Oskar sequence was identified in this species. Red: putatively duplicated Oskar sequences (sequence similarity < 80%; see Methods). Families containing *oskar* duplications are highlighted as per Figure 4.

**Supplementary Figure S5:**
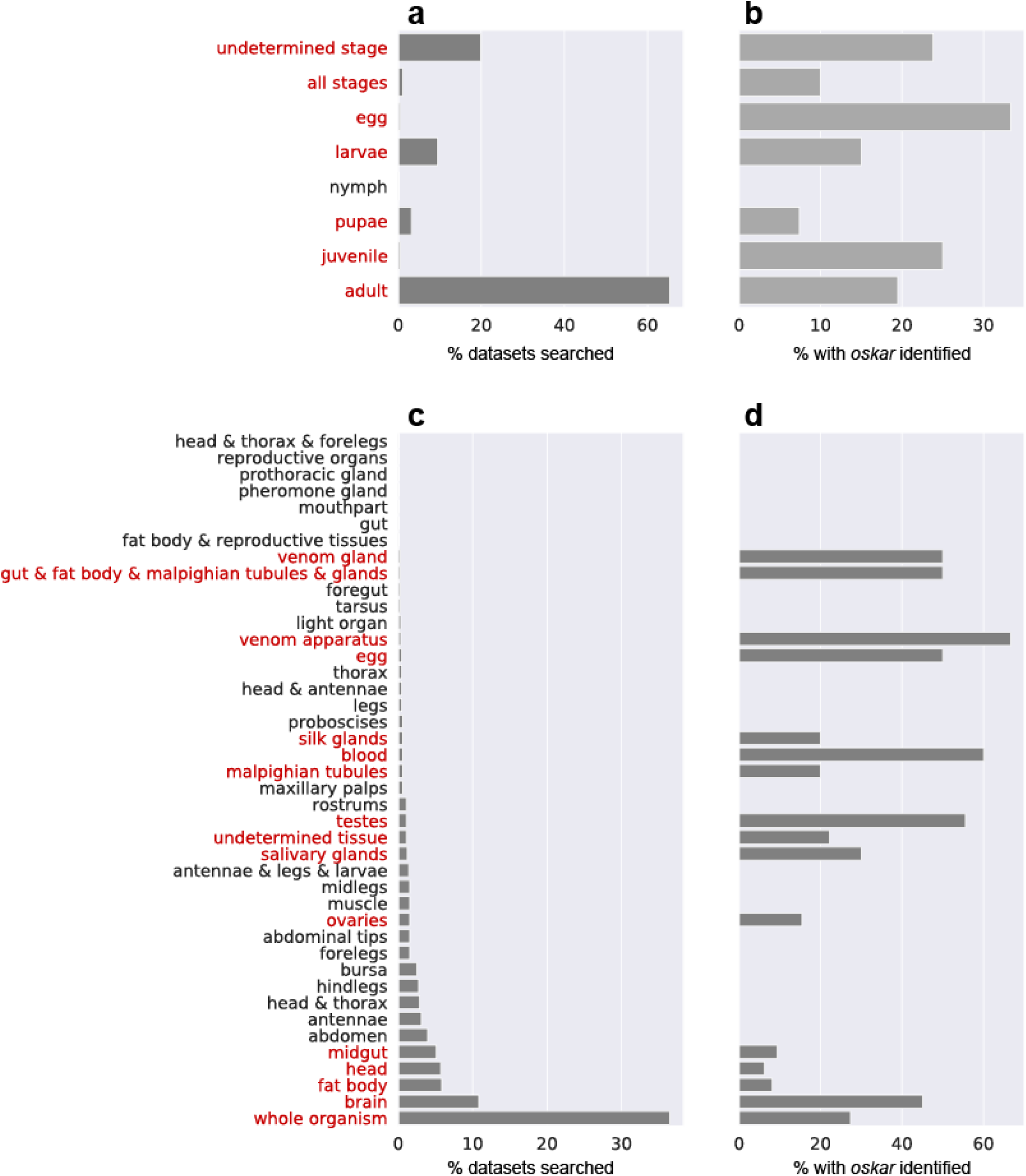
Tissue and developmental stage metadata analysis of *oskar* identification in transcriptome datasets. **(a)** Proportion of analyzed datasets that were sequenced from the developmental stages indicated on the Y axis. **(b)** Proportion of analyzed datasets per developmental stage in which an *oskar* ortholog was identified (red). **(c)** Proportion of analyzed datasets that were sequenced from the tissue type indicated on the Y axis. **(d)** Proportion of analyzed datasets per tissue type in which an *oskar* ortholog was identified (red)

**Supplementary Figure S6:**
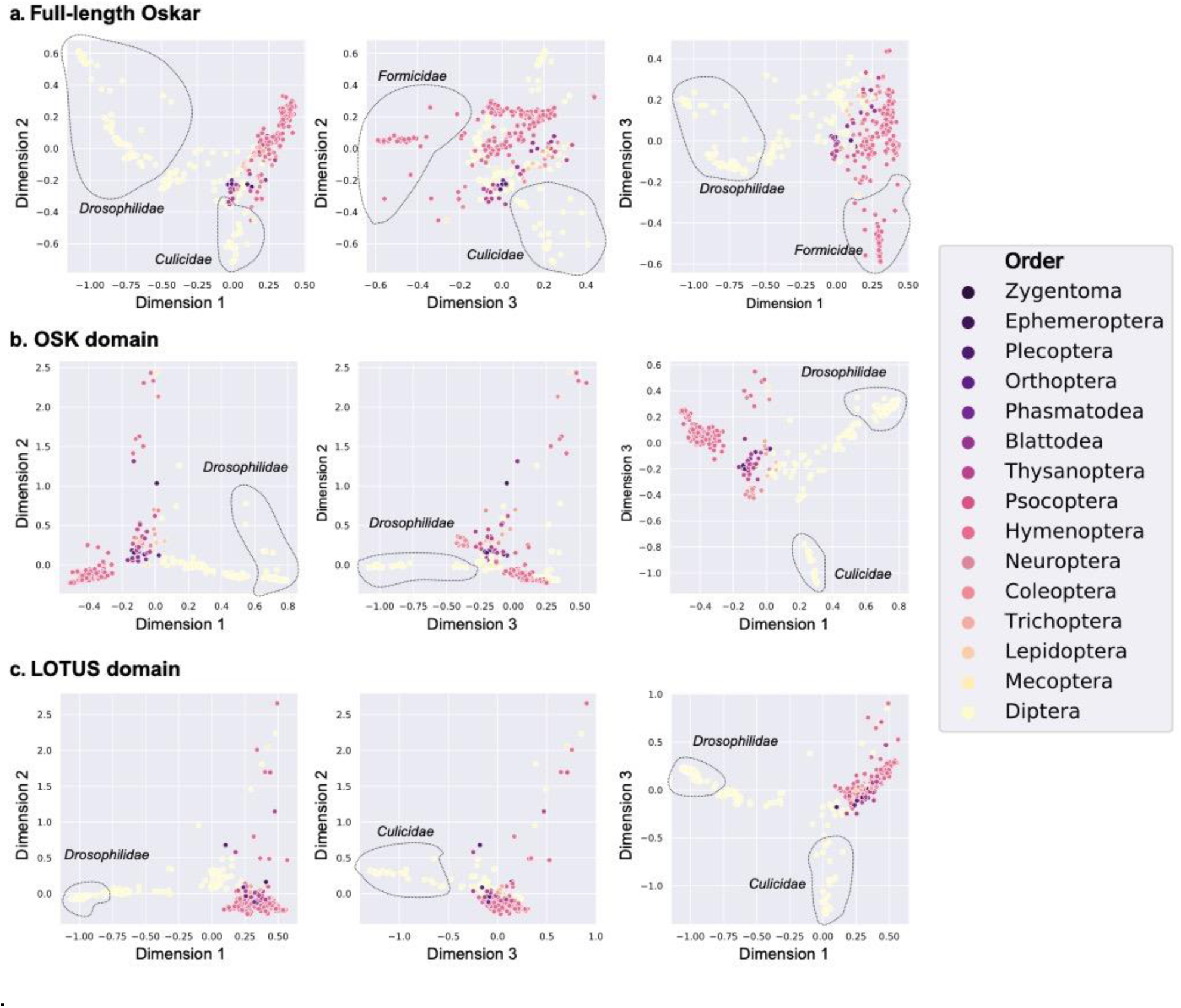
Multiple Correspondence Analysis (MCA) of full-length Oskar, the OSK domain and the LOTUS domain. MCA analysis of trimmed (30% occupancy) alignments for (a) full-length Oskar, (b) the OSK domain and (c) the LOTUS domain colored by insect order (see legend at right). The alignment was projected onto the first three main MCA dimensions (1, 2 and 3). Each dot corresponds to one sequence. Dotted line outlines specific families of interest as discussed in the text

**Supplementary Figure S7:**
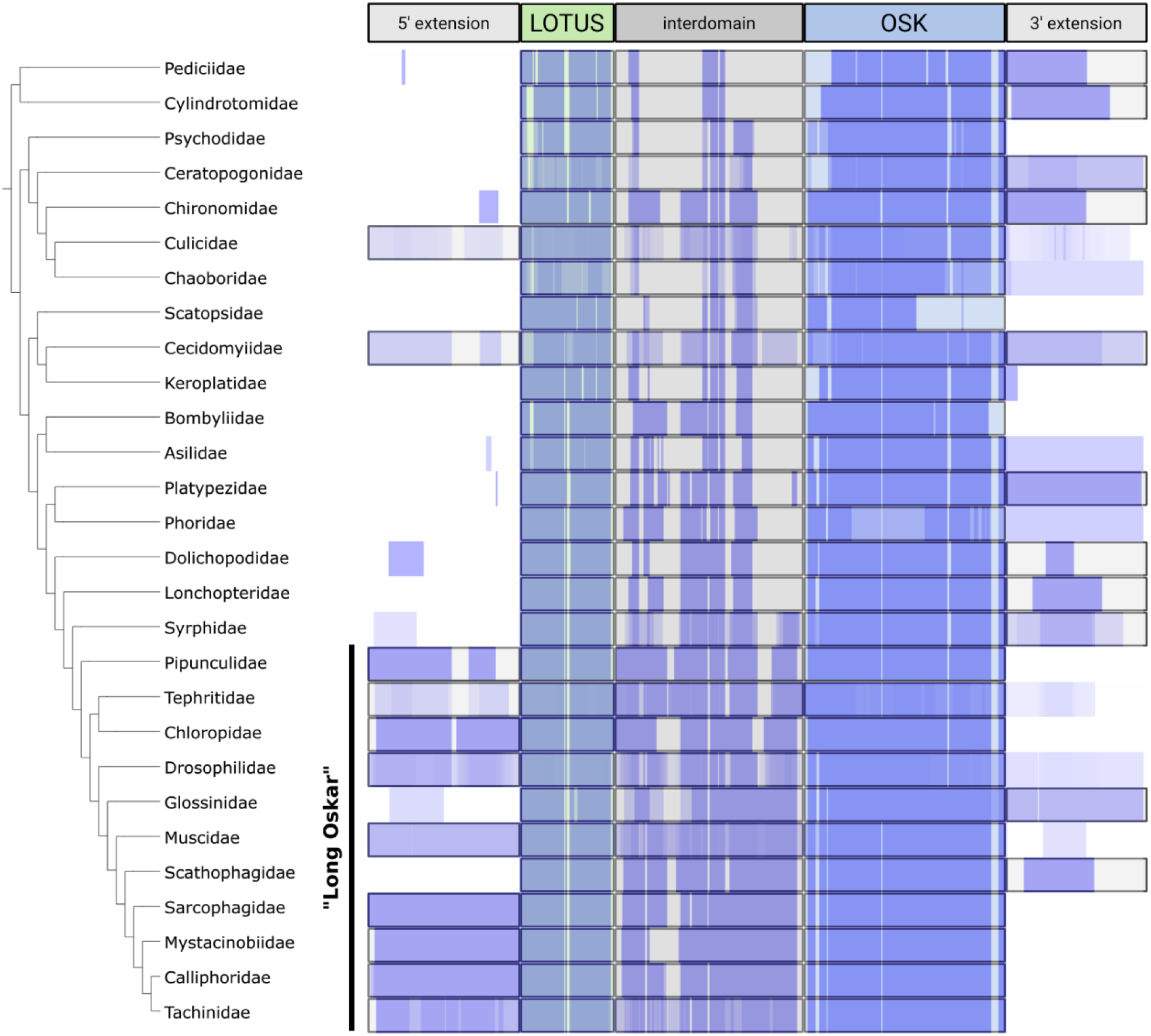
Evolution of the structure of Oskar in Diptera. Left: dipteran phylogeny from (Maddison, et al. 2007; Wiegmann, et al. 2011). Top: schematic representation of Oskar domain structure. Blue: heatmap showing the overall occupancy of an amino acid position in the Oskar alignment trimmed for at least 10% overall occupancy at a given position. For each dipteran family, occupancy at a given position is defined as (number of non-gap amino acids / number of sequences in that family). If a 3’ or 5’ extension (defined as a coding sequence unbroken by stop codons, 5’ of the first residue of the LOTUS domain, or 3’ of the last residues of the OSK domain but 5’ to a predicted poly-A tail) was detected in a family, a black box outlines the putative domain. Any such identified 5’ domains were designated as putative “Long Oskar” domains.

**Supplementary Figure S8:**
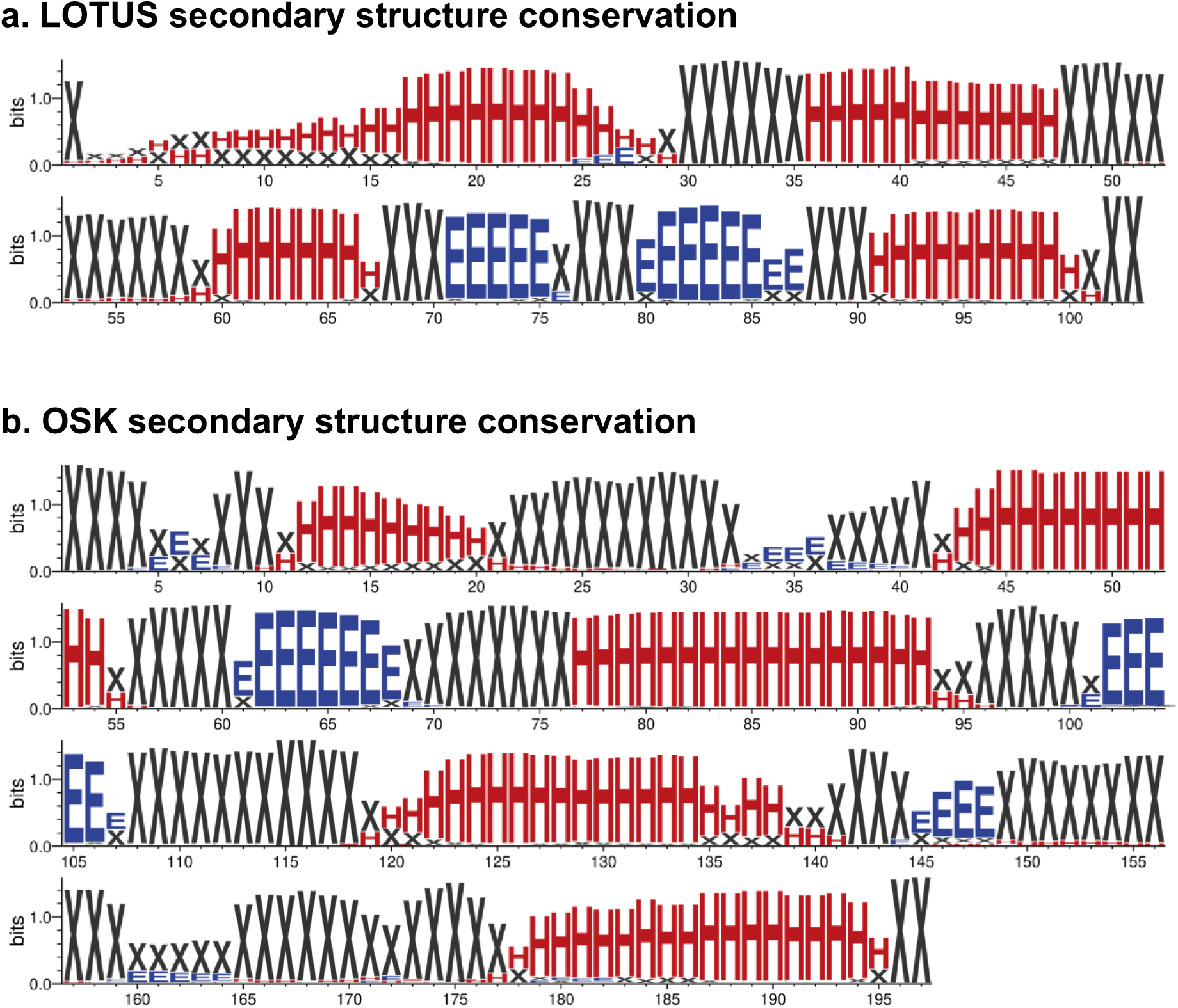
Oskar domains secondary structure conservation. Sequence Logo of Jpred4 predictions for LOTUS and OSK domains showing the conservation of secondary structures, computed with WebLogo (Crooks, et al. 2004). The height of each letter represents that state’s (X, H or B) conservation throughout the alignment in bits. X (black): unfolded amino acids; H (red): α helices; E (blue): β sheets. **(a)** Prediction for the LOTUS domain. **(b)** Prediction for the OSK domain.

**Supplementary Figure S9.**
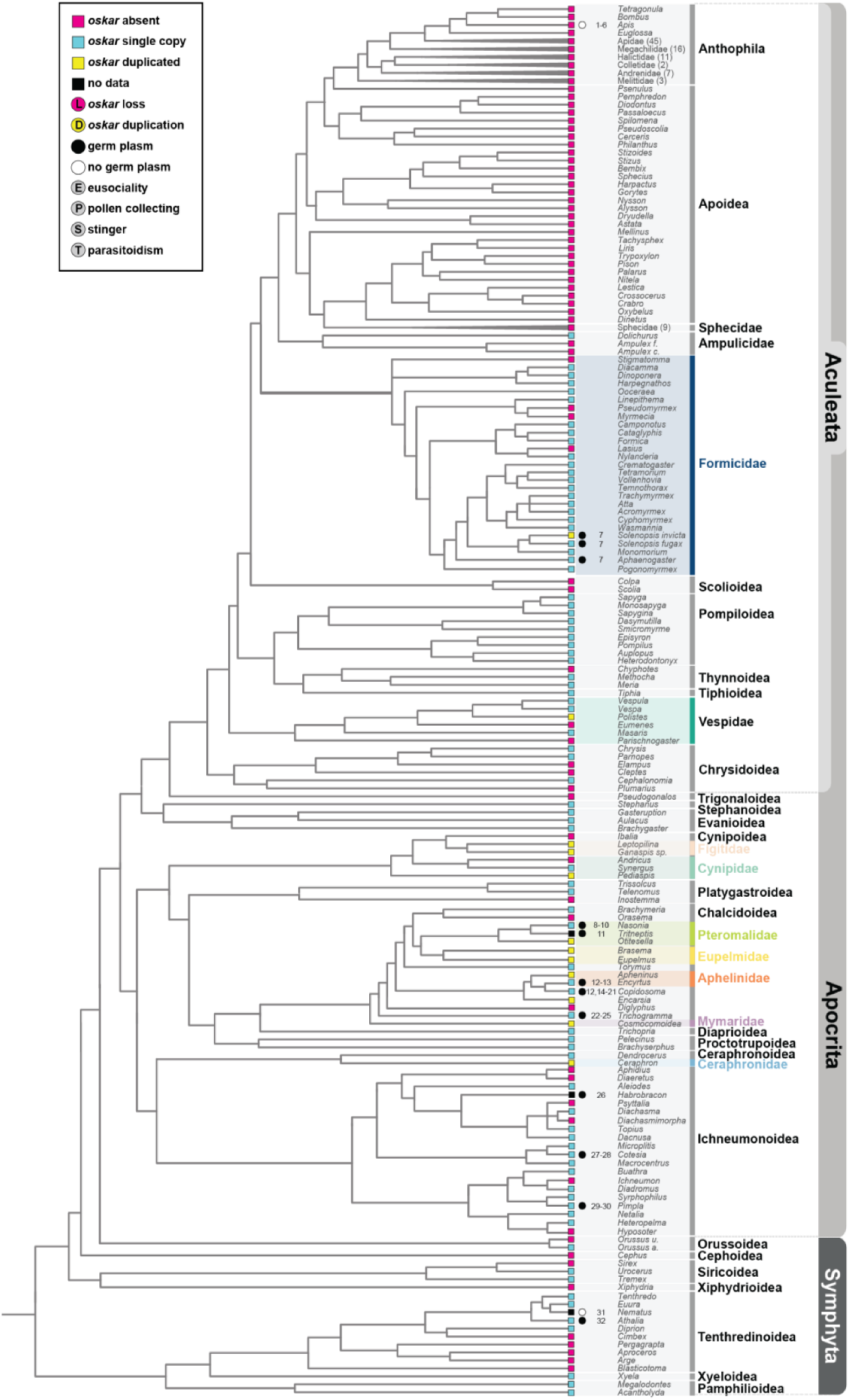
Duplications and losses of *oskar* in Hymenoptera. Absence (magenta) or presence of *oskar* orthologs detected in single copy (cyan) or multiple copies (yellow) in the genomic or transcriptomic datasets examined in this study. Genera shown in italics indicate individual species searched and are abbreviated simply for space reasons. Genera shown in regular type (not italics) indicate a summary of the results from multiple congeneric species, which were nearly always consistent within genera; in all cases where intrageneric results for *oskar* presence or absence were inconsistent, we gave precedence for the finding obtained from a genome sequence (GCF or GCA) over findings obtained from a transcriptome (TSA)), o for Hymenoptera species or genera. For some species, germ cell specification via germ plasm (black circles) or differentiation from mesoderm (no germ plasm; white circles) has been reported in the literature, with primary data references indicated by numbers as follows: [1-6]: (Bütschli 1870; Fleig and Sander 1985, 1986; Zissler 1992; Gutzeit, et al. 1993; Dearden 2006); [7]: (Khila and Abouheif 2008); [8-10]: (Bull 1982; Lynch and Desplan 2010; Lynch, et al. 2011); [11]: (Koscielska and Koscielski 1987); [12-13]: (Silvestri 1906, 1908); [12, 14-21]: (Silvestri 1906; Hegner 1914; Grbic’, et al. 1996; Strand and Grbic’ 1997; Grbic’ 2000, 2003; Donnell, et al. 2004; Zhurov, et al. 2004); [22-25]: (Gatenby 1917a; Gatenby 1917b; Gatenby 1918; Gatenby 1920); [24]: (Amy 1961); [27-28]: (Gatenby 1920; Tawfik 1957); [29-30]: (Bronskill 1959; Fleischmann 1975); [31]: (Shafiq 1954); [32]: (Sumitani, et al. 2003). Phylogenetic relationships as per (Nyman, et al. 2006; Field, et al. 2011; Schmidt 2013; Prous, et al. 2014; Ward 2014; Malm and Nyman 2015; Vilhelmsen 2015; Ward, et al. 2016; Peters, et al. 2017; Chen and Achterberg 2018; Peters, et al. 2018; Sharanowski, et al. 2021). Evolution of major hymenopteran life history characteristics (eusociality, pollen collecting, stinger, parasitoidism) as per (Peters, et al. 2017).

